# siRNA-mediated *de novo* silencing of *Ac/Ds* transposons is initiated by alternative transposition in maize

**DOI:** 10.1101/2020.01.07.897926

**Authors:** Dafang Wang, Jianbo Zhang, Tao Zuo, Damon Lisch, Meixia Zhao, Thomas Peterson

## Abstract

Although Transposable Elements (TEs) comprise a major fraction of many higher eukaryotic genomes, most TEs are silenced by host defense mechanisms. The means by which otherwise active TEs are recognized and silenced remains poorly understood. Here we analyzed two independent cases of spontaneous silencing of the active maize *Ac/Ds* transposon system. This silencing was initiated by Alternative Transposition (AT), a type of aberrant transposition event that engages the termini of two nearby separate TEs. AT during DNA replication can generate Composite Insertions (CIs) that contain inverted duplications of the transposon sequences. We show that the inverted duplications of two CIs are transcribed to produce dsRNAs that trigger the production of two distinct classes of siRNAs: a 24-nt class complementary to the TE terminal inverted repeats (TIRs) and non-coding sub-terminal regions, and a 21-22 nt class corresponding to the TE transcribed regions. Plants containing these siRNA-generating CIs exhibit decreased levels of *Ac* transcript and heritable repression of *Ac/Ds* transposition. This study documents the first case of TE silencing attributable to transposon self-initiated AT and may represent a general initiating mechanism for silencing of DNA transposons.

**Article summary:** Transposable Elements (TEs) are often silenced by their hosts, but how TEs are initially recognized for silencing remains unclear. Here we describe two independent loci that induce *de novo* heritable silencing of maize *Ac/Ds* transposons. Plants containing these loci produce dsRNA and *Ac*-homologous small interfering RNAs, and exhibit decreased levels of *Ac* transcript and heritable repression of *Ac/Ds* transposition. We show that these loci comprise inverted duplications of TE sequences generated by Alternative Transposition coupled with DNA re-replication. This study documents the first case of transposon silencing induced by AT and may represent a general initiating mechanism for TE silencing.

## Introduction

Transposable Elements (TEs) comprise a large proportion of eukaryotic genomes, including those of important crop plants such as rice (>35%, IGRS Project, Matsumoto et al. 2005), sorghum (62%) (Paterson *et al*. 2009), and maize (85%) (Schnable *et al*. 2009). Over time, multiple sporadic TE proliferations altered the number and distribution of TE sequences, enhancing genome diversity between species, and even among varieties of the same species (Tikhonov *et al*. 1999; Sanmiguel and Vitte 2009). These observations indicate that TEs have been, and continue to be, a major natural force driving genome evolution.

Although uncontrolled TEs can be deleterious to the host (Kidwell 1985), the majority of the transposons in genomes remain silenced and genome integrity is effectively preserved. Small RNA-mediated silencing is an efficient means to initiate and maintain the repression on both Class I (RNA) and Class II (DNA) TEs. For example, the Pol IV-RDR2 pathway generates 24-nt siRNAs to mediate CG, CHG and CHH methylation at the homologous DNA target. Because it relies on transcripts from methylated templates, the Pol IV-RDR2 pathway likely serves to reinforce or maintain silencing of previously-silenced TEs (reviewed by Law and Jacobsen 2010; Haag and Pikaard 2011; Castel and Martienssen 2013; Matzke et al. 2014). Although the mechanisms of maintenance of TE silencing have been extensively studied, the mechanisms responsible for *de novo* silencing remain largely obscure. The first evidence for a spontaneous silencer of an active DNA TE came from analysis of *Mu killer* (*Muk),* a locus that can heritably silence *MuDR* transposons in maize. *Muk* comprises an inverted duplication of the *MuDR* 5’ terminal inverted repeat (TIR) and a portion of the *mudrA* gene, which is required for element excision (Lisch, 2002 review). This long inverted repeat is transcribed into dsRNAs which initiate siRNA-mediated silencing of intact *MuDR* elements (Slotkin *et al*. 2003, 2005b; Li *et al*. 2010b). Recent studies of *de novo* silencing of active LTR retroelements in *Arabidopsis* demonstrated the involvement of RDR6-dependent RNA-dependent DNA Methylation (RdDM) (Marí-Ordóñez *et al*. 2013a; McCue *et al*. 2014; Duan *et al*. 2015; Panda *et al*. 2016a). RDR6 synthesizes dsRNAs from the Pol II transcript of active TEs, which is then processed into 21-22 nt siRNAs (Nuthikattu *et al*. 2013); these 21- 22 nt siRNAs can then induce Post-Transcriptional Silencing (PTGS) as well as Transcriptional Gene Silencing (TGS) when they are loaded onto AGO2 (Pontier *et al*. 2012). This complex is thought to interact with PolV and DNA and histone methyltransferases in order to methylate homologous DNA sequences (Matzke *et al*. 2014). Silencing can then be maintained via the classic RdDM pathway, which involves Pol IV-mediated transcription from previously methylated sequences.

Maize *Ac/Ds* transposable elements were the first transposons to be discovered and characterized (McClintock 1948b, 1949, 1950; McClintock 1951). As members of the Class II *hAT* transposon superfamily, *Ac/Ds* elements are less numerous than Class I retroelements, which are often highly amplified (Sanmiguel and Vitte 2009). However, *Ac/Ds* elements can strongly impact gene expression due to their preferential insertions into genes (Vollbrecht *et al*. 2010a) and the induction of genome rearrangements via Alternative Transpositions (AT) (Zhang and Peterson 1999). Unlike standard transposition reactions which act on the 5’ and 3’ termini of a single element, Alternative Transposition (AT) acts on the termini of two separate, usually nearby elements (Gray *et al*. 2000). During AT, the termini of two nearby separate TEs can interact with the transposase and insert into linked or unlinked sites, resulting in various chromosomal rearrangements, including duplications, deletions, inversions and translocations (Zhang and Peterson 2004; Huang and Dooner 2008; Zhang *et al*. 2009; Wang *et al*. 2015). Moreover, the occurrence of AT during DNA replication can generate novel structures termed Composite Insertions (CIs), which are conglomerates of transposon and host genome sequences. Due to their complex and heterogeneous structures, CIs represent an ongoing source of diverse *Ac* sequence configurations in the genome (Zhang *et al*. 2014a).

In a previous study of *Ac/Ds* AT, Zhang and Peterson (Zhang and Peterson 1999) identified an allele that induced significant repression of *Ac*/*Ds* transposition in *trans.* Because this stock contained only one copy of *Ac*, the observed *Ac/Ds* repression could not be explained by the classic *Ac* negative dosage effect, in which increased *Ac* copy number results in reduced frequency and developmental delay of *Ac/Ds* transposition (McClintock 1949; Brink, R.A., Nilan 1952). Subsequently, we isolated a second allele independently derived by AT that exhibits a similar novel *de novo* repression of *Ac/Ds.* Here we show that both of these alleles: 1) cause *de novo* and heritable repression of *Ac/Ds* transposition; 2) contain CIs with inverted duplications of *Ac* sequence; 3) produce *Ac-*homologous dsRNA transcripts driven by either a flanking host gene promoter or the *Ac* promoter; and 4) accumulate 21, 22, and 24-nt siRNAs corresponding to the region of dsRNAs transcribed from each CI. The siRNA profile includes two distinct classes: a 24-nt class corresponding to the TIR and sub-terminal region of *Ac/Ds,* and a 21-22 nt class homologous to portions of the transcribed region of *Ac*. These results confirm and extend a previous model of AT-induced DNA re-replication to generate novel CIs (Zhang *et al*. 2014a), and also show that the resulting CIs can confer a new genetic function (transposon silencing). In addition, we provide the first evidence for heritable siRNA-mediated silencing of *Ac/Ds* activity. More generally, this study is the first demonstration of *de novo* TE silencing induced by AT, which may represent a general mechanism of self-repression of Class II transposons.

## Materials and Methods

### Maize stocks and screen

In maize nomenclature, *p1* alleles can express color in the kernel pericarp (the maternal tissue surrounding the seed) and/or the cob. The first letter designates pericarp expression and the second designates cob expression: “w” for white, “r” for red, and “v” for variegated. The progenitor *p1-vv-9D9A* allele was introgressed in inbred B73 background (*p1-wr*). In order to screen for new variants in the *Ac* activity from *p1-vv-9D9A* progeny, we crossed *p1-vv-9D9A/ p1-wr* plants to the *Ac* tester line, *p1-ww; rm-3::Ds*. In the ears harvested, we screened for multi-kernel sectors or individual kernels with developmentally delayed (small) purple spots on kernel aleurone due to the repressed *Ac* activity and consider them as *p1-ww-id* candidates. Kernels of *p1-ww-id* candidates were then planted and backcrossed to B73 for more than six generations to generate the material for study. After introgression, the final genotype of plants carrying these candidates is *p1-ww-id/ p1-wr; r1/r1* in a B73 background.

### Plant growth and tissue collection for RNA isolation and small RNA sequencing

Seeds were germinated in SB 300 Universal Soil and grown in a PGW-40 growth chamber (Swanson-Wagner et al. 2006); 25° for 15 hours in the light and 20for nine hours in the dark. Both the above-ground tissues (shoot) and below-ground tissues (root) were harvested. Nine random plants were pooled for each genotype and tissue type. Total RNA was extracted by PureLink Reagent (Life Technologies) from seedling shoots and roots 14 days after sowing, then treated with DNase I (NEB) to remove residual genomic DNA. cDNA was prepared by reverse transcription primed by Oligo-dT (Thermo Scientific) as the template for PCR.

### Polymerase Chain Reaction (PCR)

Total DNA was prepared by using a modified CTAB (cetyltrimethylammonium bromide) extraction protocol (Allen *et al*. 2006) on shoot tissue from young plants. HotMaster Taq polymerase from 5PRIME, Inc. was used in the PCR reactions. PCR reactions were heated at 94° for two min; followed by 35 cycles of 94°C for 20 seconds, 60-68 ° for 30 seconds, and 65 ° for 1 min/1 kb expected product length; followed by a final cycle at 65 ° for eight minutes. The sequences of oligonucleotide primers are listed in Table S3.

### Southern blot

Total DNA extracted from seedling shoot was digested with restriction enzymes from Promega and electrophoresed through 0.8% agarose gels. Blotting and hybridization was performed according to standard protocols (Sambrook J. *et al*. 1989); blots were washed in stringent conditions (0.5% SDS, 0.5×SSC at 60 °).

### Detection of dsRNAs

Total RNA, treated with DNase I (NEB), prepared as described above was treated with RNasesA/T1 (Thermo Scientific) with concentrations of 0 U, 1.5 U and 15 U at 37 °C for 15 min. The treated RNA was precipitated and reverse transcribed into cDNA by SuperScript® III Reverse Transcriptase (Life Technologies) primed with random primers (Thermo Scientific) as the template for PCR. First round of PCR used primers a12 + a6, followed with the nested PCR using primers a4 + a6.

### qRT-PCR

qRT-PCR was performed on Stratagene Mx4000 multiplex quantitative PCR system. Total RNA was extracted by RNeasy Plant mini kit (Qiagen) from seedling shoots 14 days after sowing. cDNA was prepared by using Omniscript RT kit (Qiagen) primed with Oligo-dT on total RNA. PCR was catalyzed by SsoFast™ EvaGreen® Supermixes (Bio-rad) with two technical repeats and three biological repeats.

### Small RNA high-throughput sequencing and data analysis

Total RNA was extracted as described above and previously (Zuo et al., 2016); library preparation and sequencing on Illumina platform HiSeq 2000 were performed by Beijing Genomics Institute (BGI). Adapter sequences, contamination and low-quality reads were filtered from raw data. The small RNA sequences were mapped to *Ac* full length DNA sequence (4565 bp) by Bowtie (Langmead et al. 2009). Only perfectly matched short reads were included in the analysis. The mapped reads from each library were normalized to read counts per million reads.

### Data availability

The small RNA high-throughput sequencing data analyzed here is under accession number SRP062285 in NCBI. Maize stocks are available upon request. All data necessary for confirming the conclusions of the article are present within the article, figures, and tables. Supplemental material available at Figshare: https://figshare.com/s/98417663e12dd1055984.

## Results

### *p1-ww-id1* and *p1-p1-ww-id4* alleles induce *de novo* silencing of *Ac*

The maize *p1* gene is required for kernel pericarp (seed coat) pigmentation (Zhang and Peterson 1999). The *p1-vv-9D9A* allele contains an active *Ac* element and a *fAc* (*fractured Ac;* a terminally-deleted *Ac* fragment) in intron 2 (Figure 1A), which block *p1* expression and result in colorless pericarp. The somatic excision of *Ac* restores *p1* function and thus produces red clonal sectors on the kernel pericarp. Zhang and Peterson (1999) identified an inverted duplication allele, *p1-ww-id1,* and a corresponding deletion, *p1-ww-def1*, as the reciprocal products of Sister Chromatid Transposition (SCT), a type of AT (Figure 1F). We subsequently identified a second allele, *p1-ww-id4,* from other progenies of *p1-vv-9D9A*.

**Figure 1.**
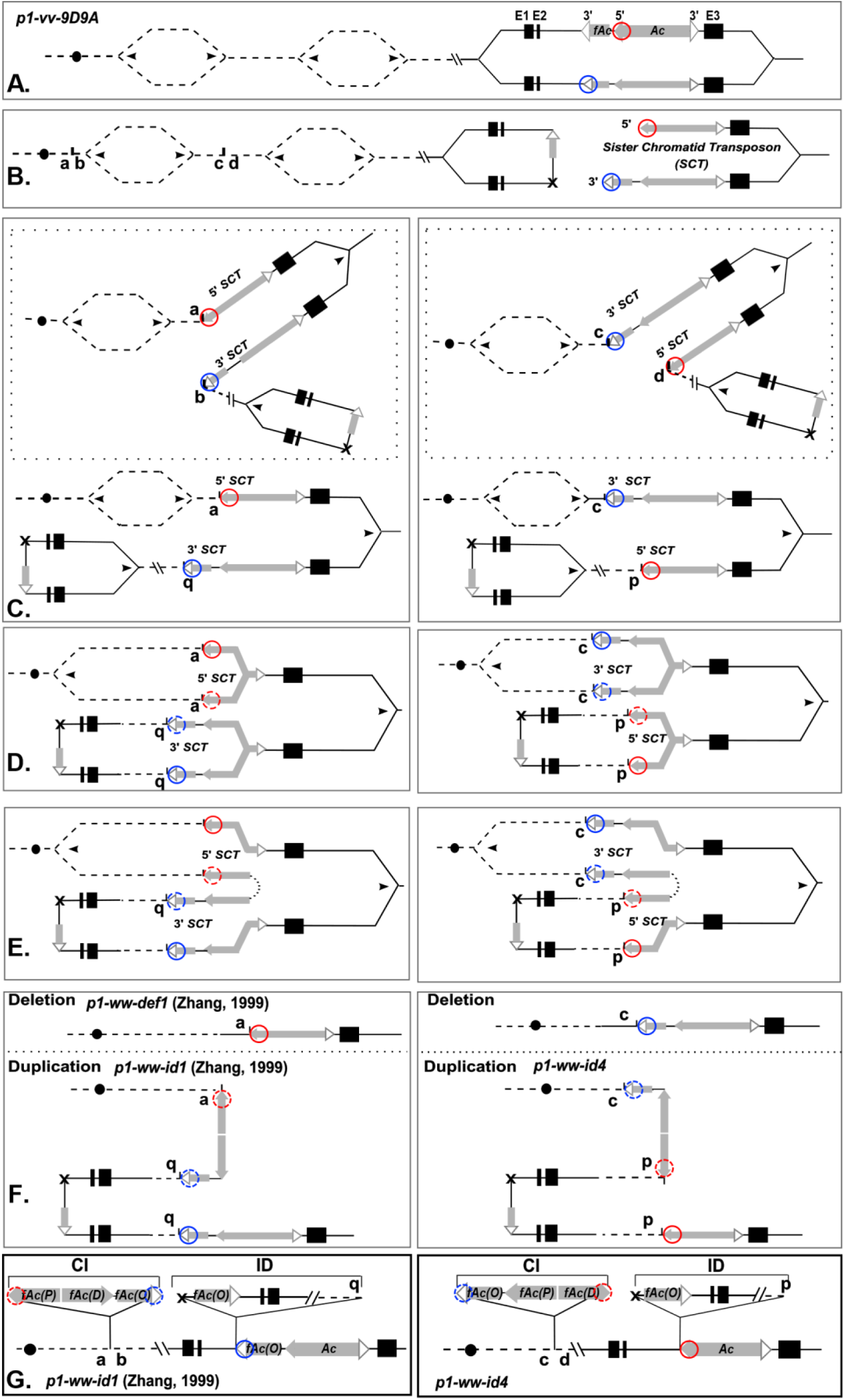
Model of Sister Chromatid Transposition (SCT)-induced DNA-re-replication. **A.** Maize chromosome 1 and progenitor allele *p1-vv-9D9A* with DNA replication bubbles. Filled circle indicates centromere, and dashed lines indicate sequences proximal to *p1*. The *p1* gene contains 3 exons (black boxes) with *Ac* and *fAc* elements located in intron 2 (grey boxes with filled/open arrowheads indicate the 5’ and 3’ *Ac/fAc* termini, respectively). The 5’ and 3’ *Ac/fAc* termini involved in SCT are circled in red and blue, respectively. **B.** SCT excision from *p1* gene results in ligation of flanking host sequences, joining two sister chromatids at site of excision footprint marked by X. Panels **C – G** show outcomes of two possible orientations of insertion into unreplicated target sites proximal to *p1* locus. Left: ligation of *Ac* 5’ terminus to the proximal side of target (“a”) and ligation of *fAc* 3’ terminus to distal side of target (“b”). Right: ligation of *fAc* 3’ terminus to the proximal side of target (“c”) and *Ac* 5’ terminus to distal side of target (“d”). **D.** Replication bubbles enlarge and re-replicate the 5’ and 3 termini of the *SCT*. **E.** Re-replication of *Ac* is aborted, releasing “broken ends” which fuse (dotted lines) to generate *Ac* inverted repeats. **F.** Chromosome DNA replication is completed, generating two sister chromatids that are segregated to daughter cells at the ensuing mitosis. One chromosome (Deletion, upper) lacks segment extending from “b” (left panel) or “d” (right panel) to *p1*, while second chromosome (Duplication, lower) contains inverted duplications of *Ac* and sequences from target sites “b” or “d” to *p1*. **G.** Schematic structures of duplication chromosomes showing Composite Insertions (CI) and Inverted Duplications (ID) as triangles inserted into the backbone of the progenitor *p1-vv-9D9A* chromosome.

As described in Zhang’s paper (Zhang and Peterson 1999), *p1-ww-id1* exhibited a decreased level of *Ac* activity. We confirmed the *Ac* repressed phenotype for both *p1-ww-id1* and *p1-ww-id4* alleles by crossing plants of *p1-ww-id1* or *p1-ww-id4* with *Ac* tester lines (genotype *p1-ww; r1-m3::Ds)*. The *Ac* tester line contains a non-autonomous *Ds* inserted in the *r1 (red1)*, a gene required for anthocyanin biosynthesis in kernel aleurone. Without an active *Ac*, the *Ds* element blocks *r1* function, resulting in colorless aleurone; however, with an active *Ac*, *Ds* can be transposed from *r1,* producing purple aleurone sectors (Kermicle 1980; Lechelt *et al*. 1989). The *p1-vv-9D9A* allele (progenitor of *p1-ww-id1* and *p1-ww-id4)*, contains an active *Ac,* and the cross between plants of *p1-vv-9D9A* and the *r1-m3::Ds Ac* tester line produced kernels exhibiting the coarsely-spotted, single-*Ac* pattern as expected (Figure 2A). In contrast, crosses of *p1-ww-id1* plant with the *Ac* tester line produce kernels with fine spotting (Figure 2C), while *p1-ww-id4* gave no spots in comparable crosses (Figure 2D). These results suggest that the *Ac* elements contained in *p1-ww-id1* and *p1-ww-id4* are repressed. Moreover, *p1-ww-id1* and *p1-ww-id4* induce *trans-*dominant repression of a normal active *Ac.* This can be seen in the kernels produced by crossing *p1-ww-id* alleles to the *p1-vv-9D9A* allele that contains a single active *Ac.* Kernels heterozygous for *p1-vv-9D9A/p1-ww-id1* show very few, late sectors (Figure 2C), while kernels produced by crossing *p1-vv-9D9A* with *p1-ww-id4* had no sectors at all (Figure 2D).

**Figure 2.**
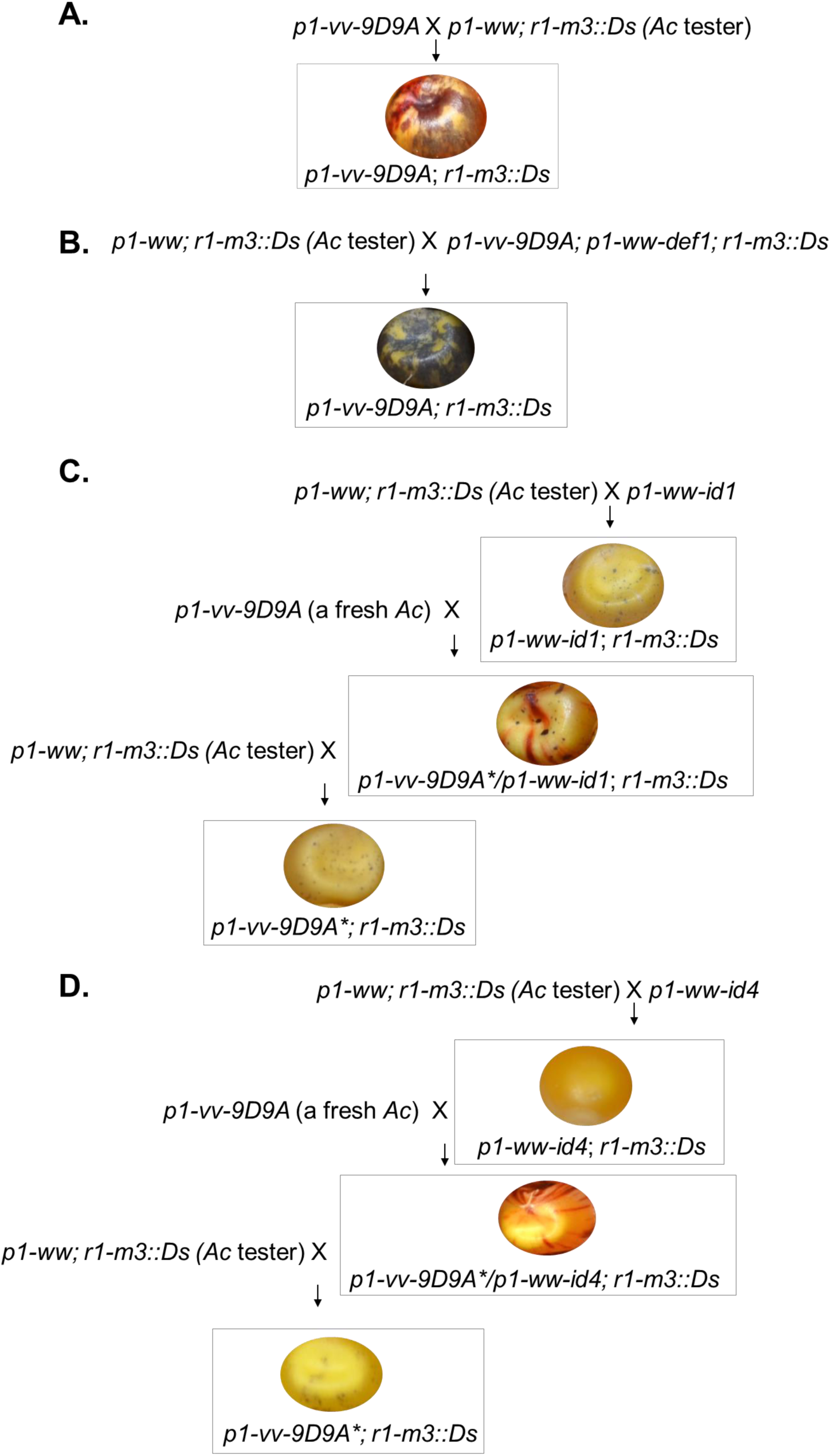
Genetic crosses indicate the *in-trans* and heritable repression of *Ac/Ds* initiated by *p1-ww-id1* and *p1-ww-id4*. **A.** *Ac* is active in the progenitor line *p1-vv-9D9A*, shown as the heavy spots on the aleurone in the cross between *p1-vv-9D9A* and *Ac* tester. **B.** The active *Ac* in *p1-vv-9D9A* cannot be repressed by another active *Ac* in *p1-ww-def1*, shown as the heavy spots on the aleurone in the cross between *Ac* tester and *p1-ww-def1*; *p1-vv-9D9A*. **C.** *Ac* activity is repressed in the line of *p1-ww-id1*, shown as the fine spots on the aleurone in the cross between *p1-ww-id1* and *Ac* tester. The repression is *in-trans*, indicated by the newly induced repression of the active *Ac* from *p1-vv-9D9A* by *p1-ww-id1.* The silencing of *Ac* in *p1-vv-9D9A* is maintained once initiated, shown as the reoccurrence of the fine spots when *p1-ww-id1* is segregated in the progeny. **D.** *Ac* activity is repressed in the line of *p1-ww-id4*, shown as no spots on the aleurone in the cross between *p1-ww-id4* and *Ac* tester. The repression is *in-trans*, indicated by the newly induced repression of the active *Ac* from *p1-vv-9D9A* by *p1-ww-id4.* The silencing of *Ac* in *p1-vv-9D9A* is maintained once initiated, shown as the reoccurrence of non-spotted phenotype when *p1-ww-id4* is segregated in the progeny.

We next tested the heritability of the *trans*-dominant repression in *Ac* activity by crossing *p1-vv-9D9A/p1-ww-id* plants to the *Ac* tester line; this cross separates the *Ac* element in *p1-vv-9D9A* from *p1-ww-id1.* If the repression of *Ac* is heritable, then most or all progeny kernels would again show few, late *Ds* excision sectors. However, if the repression of *Ac* in *p1-vv9D9A* is lifted following segregation from *p1-ww-id1,* then we would expect to see ∼50% coarsely spotted kernels (containing *p1-vv-9D9A*) and ∼50% weak or non-spotted kernels (containing *p1-ww-id1*). The results show that all the kernels produced by crossing *p1-vv-9D9A/p1-ww-id1* with the *r1-m3::Ds* tester show few or zero purple revertant sectors (Figure 2C; the ear is shown in Figure S1; the segregation numbers are shown in Table S1). This indicates that the *p1-ww-id1* allele induces *trans-*dominant, heritable silencing of *Ac* activity. Parallel genetic tests show that *p1-ww-id4* also induces silencing of *Ac,* but the heritability of repression is less stable than that induced by *p1-ww-id1*. Crossing the *r1-m3::Ds* tester by *p1-vv-9D9A/p1-ww-id4* produced 21% coarsely-spotted kernels (Figure 2D, Figure S2 and Table S1; 34 out of 160 total kernels are heavily spotted; Fisher exact test statistic value is < 0.00001). In other words, *p1-ww-id4-*induced silencing of *Ac* in *p1-vv-9D9A* was maintained in only ∼58% of progeny kernels. As a control, plants of genotype *p1-vv-9D9A*/*p1-ww-def1* were crossed by the *Ac* tester line. The *p1-ww-def1* allele carries a standard, non-repressing *Ac* in the *p1* locus; all of the progeny kernels show typical early, coarse spotting from the active *Ac* in either *p1-vv-9D9A* or *p1-ww-def1* (Figure 2B; the ear is shown in Figure S3).

To test the maintenance of silencing, plants containing a silenced *p1-vv9D9A* allele (*p1-vv-9D9A\**/*p1-ww*; *p1-vv-9D9A\** indicates silenced by *p1-ww-id1* or *p1-ww-id4*) were crossed again to the *Ac* tester stock *r1-m3::Ds.* In these crosses, 50% of the kernels will receive *p1-ww* (no *Ac*) and be non-spotted; while the remaining 50% of kernels will receive *p1-vv-9D9A\** and their spotting pattern will reflect the maintenance of *Ac* silencing. Interestingly, *p1-ww-id1* and *p1-ww-id4* induced different levels of silencing maintenance. For *p1-vv-9D9A\** silenced by *p1-ww-id1* and separated from it for two generations, all the kernels are non-spotted or weakly spotted, indicating near complete maintenance of *Ac* repression (Figure S1 and Table S1). In contrast, *p1-vv-9D9A\** silenced by *p1-ww-id4* resulted in 36% of kernels with high frequency of *Ds* excision (Figure S2 and Table S1; 61/170 kernels are heavily spotted; Fisher exact test statistic value is 0.0116). These results indicate that *Ac* repression is maintained in only ∼28% of kernels containing *p1-vv-9D9A\** silenced by *p1-ww-id4* and separated from it for two generations.

To determine the mechanism of *Ac* repression, we performed qRT-PCR experiment using *Ac-*specific primers. The *p1-ww-id1* and *p1-ww-id4* alleles express significantly decreased levels of *Ac* transcript compared to the progenitor allele *p1-vv-9D9A* (student t-test p-value=0.0015 and 0.0004, respectively) (Figure 3), suggesting that repression occurs via transcriptional and/or post transcriptional gene silencing (TGS or PTGS).

**Figure 3.**
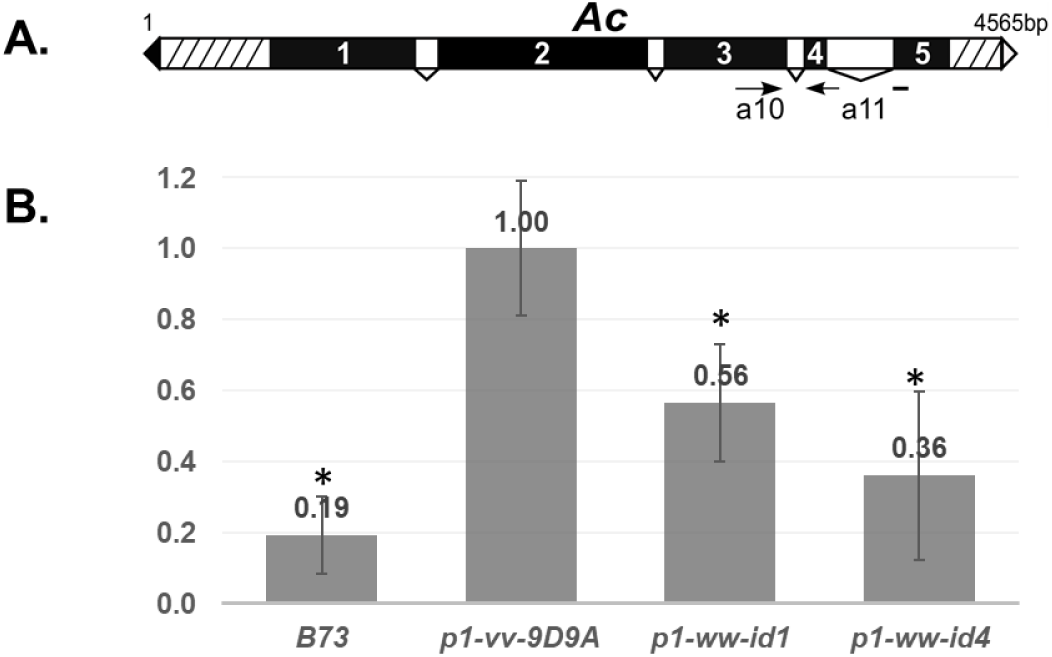
The *Ac* transcript level is decreased in *p1-ww-id* alleles. **A.** Schematic structure of the 4565 bp *Ac* transposable element. Filled and open triangles indicate *Ac* 5’ and 3’ terminal inverted repeats (TIR), respectively. Black boxes with numbers are *Ac* exons 1 – 5; white boxes with subtending lines are introns; and hatched boxes label the *Ac* subterminal regions. Primers used in the q-RT PCR (a10 and a11) are labeled as arrows; note that primer a11 spans *Ac* intron 4. **B.** Quantitative RT-PCR measurement of *Ac* transcript levels. Transcript levels in each allele are normalized to that of *p1-vv-9D9A*. Bars labeled by a star indicate transcript levels that are significantly different from *p1-vv-9D9A* by the student’s t-test with critical value at 0.05.

### Silencing alleles contain Inverted Duplications (IDs) and Composite Insertions (CIs) that are predicted by the SCT-induced DNA re-replication model

We previously reported that tandem direct duplications and associated CIs can be produced by RET (Reversed End Transcription), a type of AT, followed by DNA re-replication and repair (Zhang and Peterson 2004; Zhang *et al*. 2014a). The CIs are bordered by partial or full copies of the *Ac* transposon and may also include sequences flanking the original *Ac* donor site. Extending the principle of AT-induced DNA re-replication, we propose that the *p1-ww-id1* and *p1-ww-id4* alleles were generated by SCT, followed by DNA re-replication and repair to produce CIs capable of silencing the *Ac* transposon system (Figure 1 and Video 1). In SCT, the *Ac* transposase acts on a pair of directly-oriented *Ac* termini that are present in the progenitor allele *p1-vv-9D9A* (Figure 1A). This allele contains a complete *Ac* element inserted 112 basepairs from a second element termed *fractured Ac* (*fAc*), which contains only the 3’ half of *Ac.* Importantly, the 5’ TIR of *Ac* and the 3’ TIR of *fAc* are oriented in the same direction. The *Ac* element is known to preferentially transpose during or shortly after DNA replication in the cell cycle (Greenblatt and Brink 1962; Chen *et al*. 1987), possibly because the *Ac* transposase preferentially interacts with hemi-methylated *Ac* TIRs (Ros and Kunze 2001). Following replication, the *Ac* 5’ and *fAc* 3’ termini located on sister chromatids have strand-specific hemi-methylation patterns that are competent for transposition. Excision of the 5’ and 3’ TIRs followed by religation of the host sequences flanking these termini generates a sister chromatid fusion with a small sequence footprint at the excision site (Weil and Wessler 1993) (Figure 1B). The excised TIRs may then reinsert at many possible genomic sites; in the case of *p1-ww-id1* and *p1-ww-id4,* the termini inserted into proximal, unreplicated target sites (Figure 1C-D). As shown previously, SCT followed by insertion into a proximal site generates reciprocal duplication/deletion chromatids (Zhang and Peterson 1999). Insertion of the *Ac* termini into the target site generates 8-bp target site duplications (TSD), a diagnostic feature of *Ac* integration. Following insertion, DNA replication fork progression continues into the newly-inserted *Ac* termini, re-replicating the *Ac*/*fAc* sequences until the DNA re-replication forks spontaneously abort. In some cases, re-replication may extend for 10 kbp or more (Zhang *et al*. 2014b); for *p1-ww-id1* and *p1-ww-id4*, we propose that re-replication aborts before completing the replication of the full length *Ac*. The abortion of replication forks produces two broken ends that fuse and generate two *fractured Ac (fAc)* in inverted orientation (Figure 1E). We term the proximal one *fAc(P)*, and the distal one *fAc(D),* to distinguish them from the original *fAc(O)* in the progenitor *p1-vv-9D9A* allele. Following replication and repair, the two sister chromatids (one with a duplication, and one with a corresponding deletion) segregate at the ensuing anaphase of mitosis into two daughter cells (Figure 1F). Importantly, the two new *fAcs* generated by SCT are both derived from *Ac* 5’ termini and are in an inverted orientation relative to each other (Figure 1G).

To test this model, we first examined the presence of the predicted diagnostic sequence features on the *SCT* excision. The excision is expected to leave sequence footprints in *p1* and at the center of the duplicated segments, as previously reported for *p1-ww-id1* (Zhang and Peterson 1999). For *p1-ww-id4,* we PCR amplified and sequenced this region (using primers a3 + P1(fAc) in Figure 4C; results shown in Figure 4D). Similar to the footprint of *p1-ww-id1* (Zhang and Peterson 1999), *p1-ww-id4* shows typical changes on the first nucleotide at each side of the flanking sequence (File S1).

**Figure 4.**
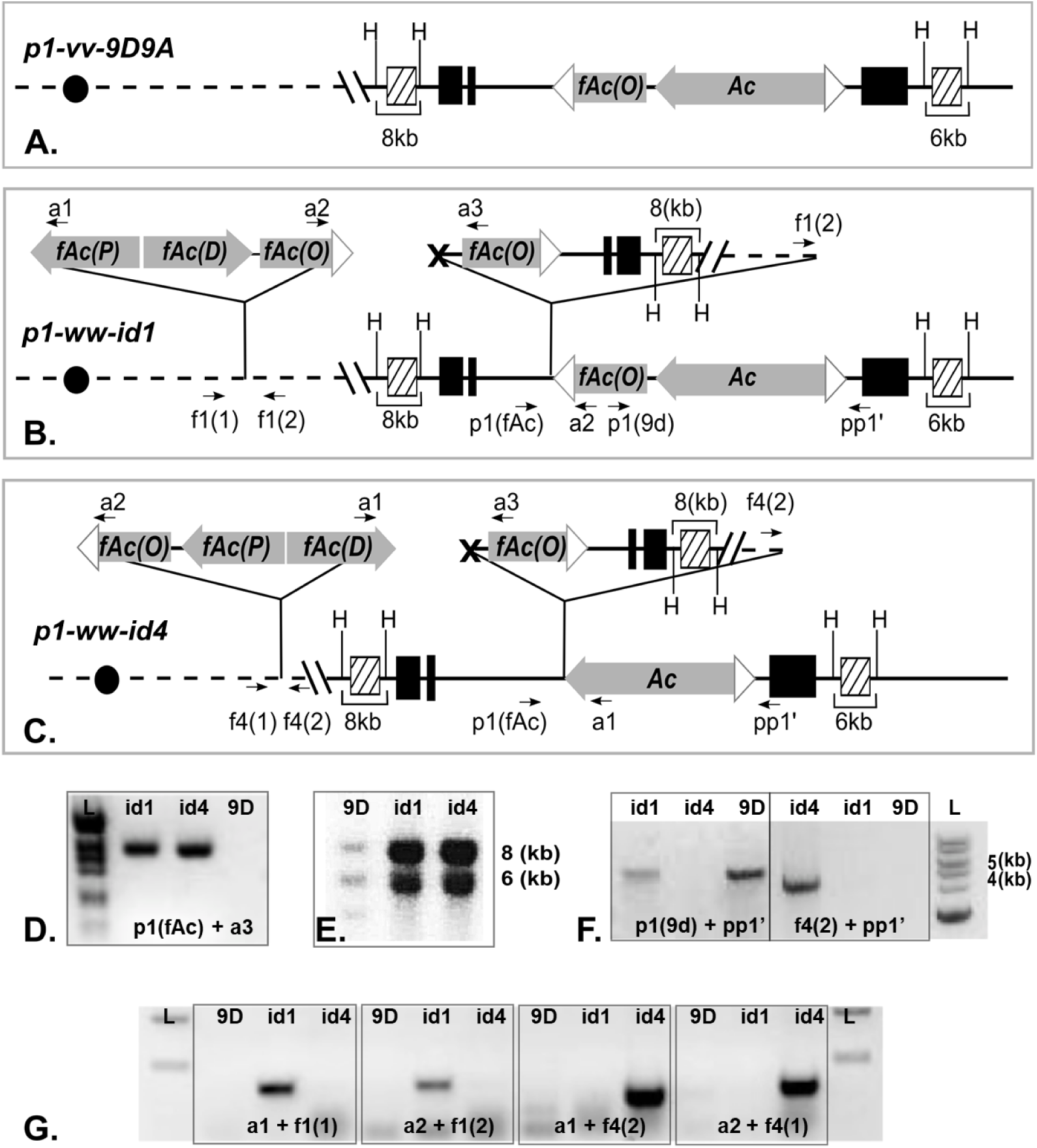
Structure of Inverted Duplications and Composite Insertions in *p1-ww-id1* and *p1-ww-id4*. **A-C.** Schematic structures of the progenitor allele *p1-vv-9D9A,* inverted duplication alleles *p1-ww-id1* and *p1-ww-id4*. PCR primers are labeled as arrows, and sequences homologous to hybridization probe *15* are labeled as hatched boxes. Other symbols as in Figure 1. **D.** PCR amplification of the *SCT* excision footprints in *p1-ww-id1* and *p1-ww-id4*. Bands were excised from the gel and sequenced (File S1). **E.** Genomic Southern blot produced by digestion with *Hind*III digestion and hybridization to probe *15*. In the progenitor allele *p1-vv-9D9A* (labeled as 9d), the intensities of the 8 and 6 kb bands are approximately equal, consistent with 1 copy of each fragment in *p1-vv-9D9A* (part A). In the *p1-ww-id1* and *p1-ww-id4* alleles, the 8 kb band intensity is approximately twice that of the 6 kb band, consistent with a duplication of the segment distal to *p1* in those alleles (parts B and C). Hybridization signals in lane 9d are overall weaker due to less DNA loaded on the gel. **F.** Gel analysis of PCRs to determine the orientation of *SCT* insertion. In *p1-ww-id1* (lane 2), primers P1(9d) and pp1’ produce a band of the same size (6.6 kb) as observed in the progenitor allele *p1-vv-9D9A*, indicating that *fAc(O)* is linked to *p1* exon 3, consistent with ligation of 3’ *SCT* to the distal side of target site. In *p1-ww-id4* (lane 3), primers f4(2) and pp1’ produce a 4.5 kb band indicating that *Ac* is linked to *p1* exon 3, consistent with ligation of 5’ *SCT* to the distal side of target site. **G.** PCR amplification of the junctions of CI with target site flanking sequences. Bands were excised from the gel and sequenced to identify the target site duplications (TSD) flanking each CI (File S2).

We then isolated the CI insertion targets of SCT for both *p1-ww-id1* and *p1-ww-id4* alleles by inverse PCR. The CI insertion sites of *p1-ww-id1* and *p1-ww-id4* are mapped to positions 51.8 Mb and 48.9 Mb, respectively, on Chromosome 1, B73 RefGen_v4 reference genome. As the *p1* locus is positioned at 48.6 Mb of Chromosome 1, the inverted duplications resulted from the *SCT* re-insertion in the two alleles can be estimated as 3.2 Mb for *p1-ww-id1* and 0.3 Mb for *p1-ww-id4*. The insertion targets were both confirmed by using the primer f1(1) with a1 for *p1-ww-id1* and f4(2) with a2 for *p1-ww-id4* (Figure 4G). The comparison of sequences flanking the CIs revealed the perfect 8-bp target site duplications (TSDs) (“GCCTCGCT” in *p1-ww-id1* and “GCCCGGAT” in *p1-ww-id4*; File S2) characteristic of *Ac* transposition. The presence of duplicated segments from the SCT re-insertion was further tested by Southern blot using *Hind*III digests and probe fragment *15* (Figure 4E). The band included in the duplicated segments (8 kb) is much brighter than the band excluded from the duplication (6.6 kb) in both *p1-ww-id1* and *p1-ww-id4*, confirming the presence of a duplication extending to the proximal side of *p1* as predicted by the SCT model. In addition, *p1-ww-id1* and *p1-ww-id4* also represent two orientations of insertion that are predicted by the model. We obtained PCR bands using primers p1(9d) and pp1’ from *p1-ww-id1* but not *p1-ww-id4* (Figure 4F), indicating that *fAc(O)* flanks the ID (Inverted Duplication) segment in *p1-ww-id1*, as predicted by the model when the 5’ terminus of *Ac* ligates to the proximal side of the insertion target (“a” in Figure 1C). We also obtained a PCR band of 4.5 kb using primers f4(2) + pp1’ in *p1-ww-id4* but not in *p1-ww-id1*, indicating that *Ac(O)* (the original *Ac* in *p1-vv9D9A)* flanks the ID segment in *p1-ww-id4* as predicted by the model when the 5’ terminus of *Ac* ligates to the distal side of the insertion target (“d” in Figure 1C). In summary, these results show that the *p1-ww-id1* and *p1-ww-id4* alleles contain transposon donor excision footprints, proximal duplications, and CIs with flanking TSDs as predicted by SCT with insertion into unreplicated target sites. Together these observations indicate that SCT exhibits the characteristic features of standard *Ac/Ds* transposition.

### Composite Insertions contain inverted repeats of *fractured Ac* fragments

We performed Southern blots to determine the structures of the CIs in *p1-ww-id1* and *p1-ww-id4* (hereinafter termed CI-*id1* and CI-*id4*). These data are summarized in Figure 5A, and the raw data are presented in Figures S4. First, the total sizes of CIs were estimated from digests using enzymes that do not cut within the *Ac* element; using enzyme *Sal*I with probe *1D*, CI*-id1* is estimated to be in the range of 6.2 – 8.2 kb. Consistent with this, enzyme *Pst*I and probe *1P* give an estimated size of ∼6.7 kb. We then determined the internal structures of each CI by digesting with a series of endonucleases with known sites in the *Ac* sequences. We obtained the expected sized bands from digestion at the *Ssp*I and *Sca*I sites in *fAc(P),* and the expected sized bands from digestion at the *Ssp*I and *Bgl*I sites of *fAc(D)*. These results indicate that CI-*id1* includes the first 1842 bp sequences from the *Ac* 5’ terminus. In contrast, the band predicted from digestion with *EcoR*I and hybridization with probe *1P* (13 kb) was absent, and instead we observed a larger band. This indicates that both *fAc(P)* and *fAc(D)* are truncated prior to reaching the *EcoR*I site at position 2486 bp from the *Ac* 5’ end, and thus the total size of CI*-id1* is < 7.1 kb, consistent with the ∼6.7 kb estimate obtained from *Pst*I digestion.

**Figure 5.**
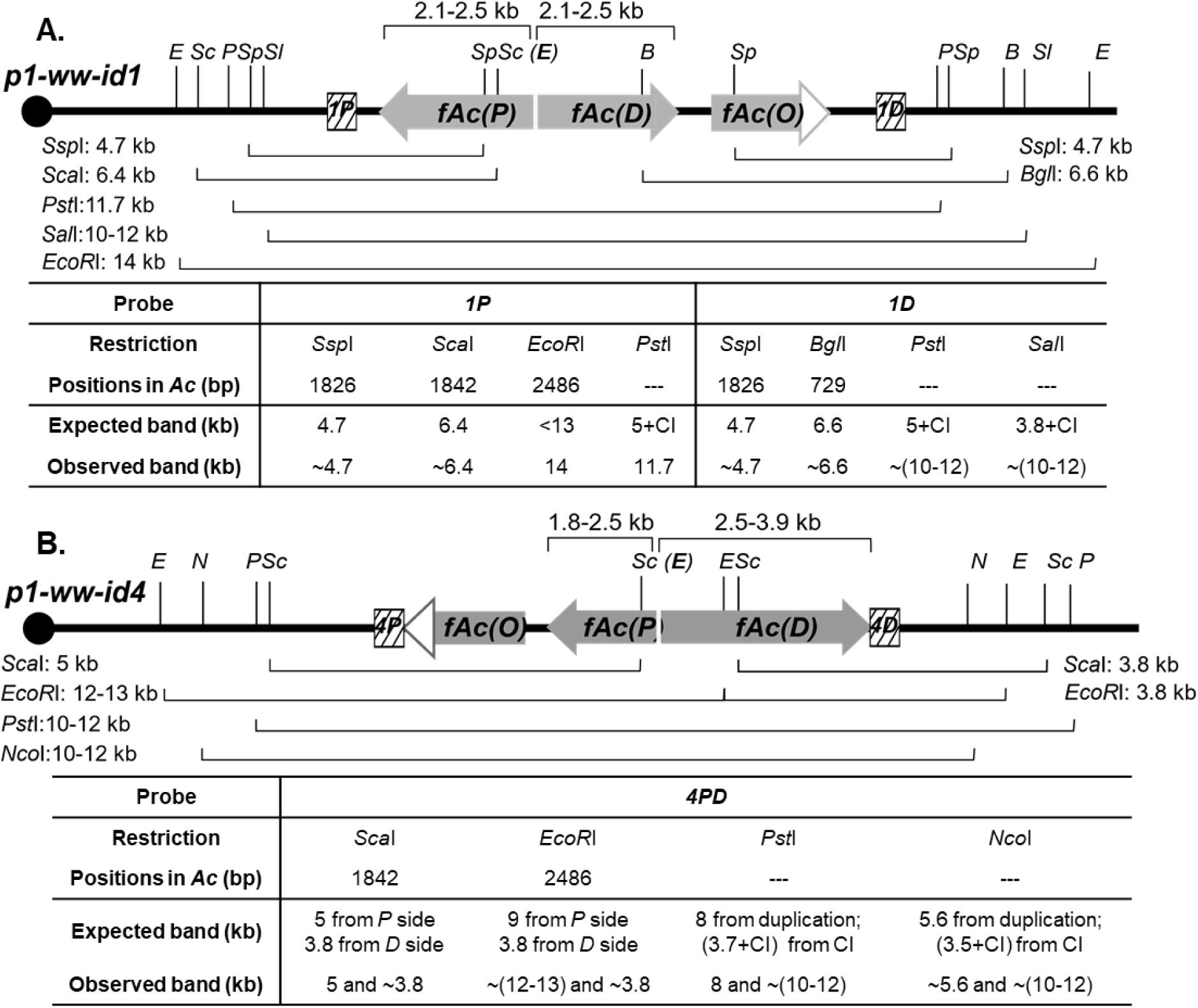
Summary of Southern blot results that elucidate the internal structures of Composite Insertions. Schematic structures of *p1-ww-id1* and *p1-ww-id4* are shown in **A** and **B**, respectively, with restriction sites (vertical lines; E: *EcoR*I; Sc: *Sca*I; P:*Pst*I; Sp: *Ssp*I; Sl: *Sal*I; B: *Bgl*I; N: *Nco*I) and probes (hatched boxes) shown. Restriction fragments that correspond to the bands from Southern blot data (Figures S4 and S5) are indicated below each allele, with their expected and actual sizes listed in the table.

Similar assays were performed on *p1-ww-id4* (Figure 5B with the raw data presented in Figures S5). *Pst*I and *Nco*I digestions both show the total size of CI-*id4* to be from 6.4-8.4 kb. CI*-id4* contains the *Sca*I site from *fAc(P)* and *fAc(D),* indicating that both *fAc* fragments include at least 1842 bp of *Ac* 5’ terminal sequences. Additionally, *fAc(D)* contains the *EcoR*I site, but *fAc(P)* does not, indicating that *fAc(P)* is truncated prior to the *EcoR*I site at position 2486 bp from *Ac* 5’ terminus. Combining the results from *Sca*I and *EcoR*I digests, the size of *fAc(P)* ranges from 1842 to 2486 bp. By subtracting the size of *fAc(P)* (1.8-2.5kb) from the total size of CI-*id4* (6.4 – 8.4 kb), we infer the size of *fAc(D)* is 2.5-3.9 kb. Extensive attempts were made to PCR amplify and sequence the exact fusion point connecting *fAc(D)* and *fAc(P)* in both CI-*id1* and CI-*id4*; unfortunately, these attempts failed, most likely due to the long perfect inverted repeats present in both CIs which would give rise to self-annealed hairpin with extensive secondary structures in the template DNA.

### The inverted repeats of *fractured Ac* fragments are transcribed by the *Ac* or flanking gene promoter to produce dsRNAs

The inverted *fAc(P)* and *fAc(D)* fragments in both *p1-ww-id1* and *p1-ww-id4* retain the *Ac* promoter. Moreover, consistent with the known preferential insertion of *Ac/Ds* into linked genes (Vollbrecht *et al*. 2010b), both CIs are located in the middle of annotated maize genes: the CI in *p1-ww-id1* is in intron 1 of Zm00001d028930, while the CI in *p1-ww-id4* is in intron 5 of Zm00001d028863 (Figure 6A-B).

**Figure 6.**
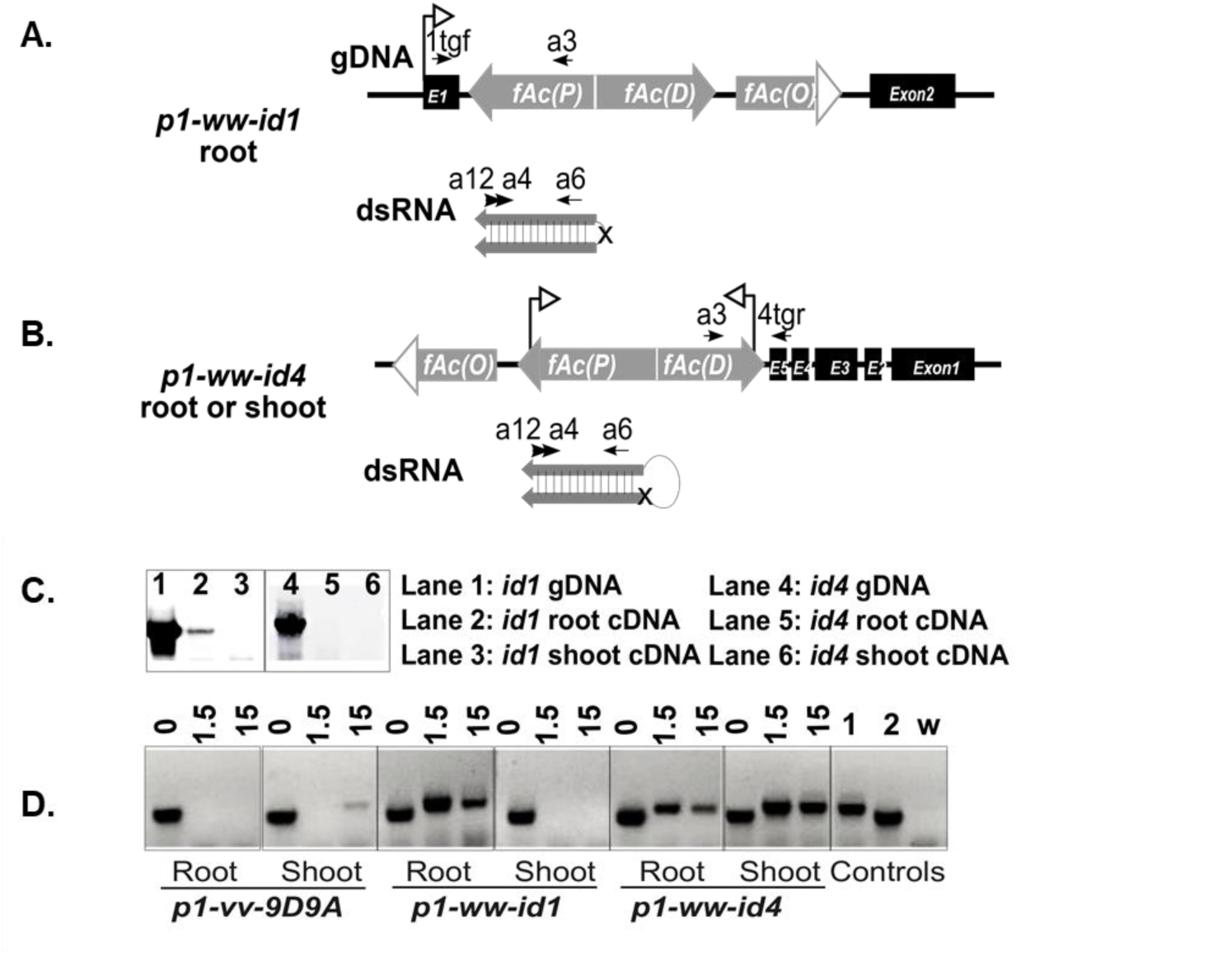
CIs in *p1-ww-id1* and *p1-ww-id4* produce dsRNA transcripts. **A.** Genomic DNA structure and dsRNA transcripts of CI in *p1-ww-id1*. The CI of *p1-ww-id1* is inserted in intron 1 of gene Zm00001d028930, and thus may be transcribed from the promoter of this host gene (arrow). The endogenous Zm00001d028930 gene is expressed in root but not shoot tissues, and the chimeric transcript and dsRNA of *p1-ww-id1* were detected only in root (Figure 6C and 6D). The absence of dsRNA in shoot tissues indicates that the *Ac* promoters are apparently not active in shoot tissues in this allele. Labeled arrows indicate the primers used in the RT-PCR. **B.** Genomic DNA structure and dsRNA transcripts of CI in *p1-ww-id4*. The CI of *p1-ww-id4* is in intron 5 of Zm00001d028863. No transcripts initiating from the promoter of this gene were detected in root or shoot (Figure 6C), suggesting that the dsRNA in this allele (Figure 6D) is produced from the endogenous *Ac* promoters (arrows). **C.** Analysis of chimeric transcripts by RT-PCR using primers 1tgf + a3 in *p1-ww-id1* (Lanes 1-3), and 4tgr + a3 in *p1-ww-id4* (Lanes 4-6). Chimeric transcript initiating from flanking gene promoter was detected only in the root of *p1-ww-id1*. **D.** Detection of dsRNA by RNAse protection assay. Total RNA from the indicated tissues and alleles were treated with DNase1 and three quantities of RNase A/T1 (“0”, “1.5” and “15” Units). Treated RNA samples were then analyzed by semi-nested PCR (primers a12+a6; followed by a4+a6) and run on gels. Bands derived from protected dsRNA are observed in *p1-ww-id1* root, and *p1-ww-id4* shoot and root, but not in *p1-vv-9D9A* (a weak band in *p1-vv-9D9A* shoot/15 may be from a loading contamination). Control lanes “1” and “2” are from templates of genomic DNA and root cDNA of *p1-ww-id1*; lane “w” is from a water template control for PCR contamination. Interestingly, the bands from protected dsRNA in the CI alleles are larger than the corresponding spliced RNA (control “2”), suggesting that the intron 1 of host gene Zm00001d028930 and the intron 1 of *Ac* are retained in the dsRNA transcript. Additional evidence for retention of introns in *p1-ww-id1* is also shown in Figure S9 and File S3.

The insertions of CIs within genes could allow their expression by read-through transcription from the flanking genes’ promoters. To test this, we designed experiments to detect chimeric gene-CI transcripts in root and shoot tissues. For *p1-ww-id1,* chimeric transcripts were detected in root, but not shoot (Figure 6); this is the same pattern of expression as for the host gene Zm00001d028930 (Wang *et al*. 2009) (Figure S6). Additionally, we detected *Ac-*homologous double-stranded RNAs (dsRNAs) in roots but not shoots of *p1-ww-id1* plants (Figure 6D). These results suggest that CI-*id1* may be transcribed from the flanking gene promoter. In contrast, gene Zm00001d028863 is not expressed in root or shoot, and we did not detect chimeric *p1-ww-id4* transcripts in those tissues (Figure 6C, File S3). However, we did detect *Ac-*homologous dsRNAs from both root and shoot, suggesting that *p1-ww-id4* is transcribed from its own *Ac* promoter. Interestingly, both the chimeric transcript and the dsRNAs seem to retain introns (File S3), suggesting that the hairpin nature of these transcripts interferes with normal splicing.

We conclude that the inverted repeats of *fractured Ac* fragments in *p1-ww-id1* and *p1-ww-id4* are transcribed into dsRNAs, either by the flanking gene promoter as in *p1-ww-id1*, or by the retained *Ac* promoter as suggested for *p1-ww-id4*. The accumulation of dsRNAs can thus provide a potential template for production of siRNAs that may directly induce *Ac* silencing.

### siRNAs derived from CI are detected in both *p1-ww-id1* and *p1-ww-id4* alleles

Encouraged by the presence of dsRNAs in *p1-ww-id1* and *p1-ww-id4*, we performed high-throughput sequencing of small RNA from root and shoot tissues of *p1-ww-id1* and *p1-ww-id4* plants; progenitor allele *p1-vv-9D9A* and standard maize inbred B73 were used as controls. As shown in Figure 7, siRNAs of 21, 22, and 24-nt that are mapped to *Ac* are dramatically enriched in both shoot and root from both *p1-ww-id1* and *p1-ww-id4* plants compared to the same tissues in *p1-vv-9D9A* and B73. These siRNAs are matched to the stem-loop region predicted from CI structures of both *p1-ww-id1* and *p1-ww-id4* (Figure 8 and Figure S7), confirming their origin from these CI transcripts. The different size classes of siRNAs map to distinctly different regions of *Ac.* The 24-nt siRNAs are enriched in the 5’ and 3’ TIR/Sub-TIR regions of *Ac* (Figure S8-B, D and E); this region includes the native *Ac* promoter and is consistent with Pol IV-RDR2 maintenance of heritable silencing. In contrast, the 21-22 nt siRNAs predominate in the transcribed regions of *Ac* (Figure S8-C). These siRNAs are associated with the PolII-RDR6 pathway and thus may induce *de novo* silencing of active *Ac* elements. Interestingly, the 21-22 nt siRNAs appear to spread from the stem to the loop region in *p1-ww-id4* (the expanded loop region is shown in Figure S7). This is consistent with a transitive process, by which RDR6 converts the single stranded RNA in the loop into double-stranded RNA, followed by the processing of dsRNAs into small RNAs. The ratio of 21 to 22 nt is increased in the loop region in shoot compared to root tissues (Table S2: chi-squared test gives p < 0.01 in shoot and p =0.24 in root), suggesting tissue-specific variation in production or stability of siRNA species. Finally, we note that 21-nt siRNAs in *p1-ww-id1* are enriched in *Ac* intron sequences, consistent with the observation that both the chimeric *Ac* transcript and dsRNAs retain introns in *p1-ww-id1* (Figure S9).

**Figure 7.**
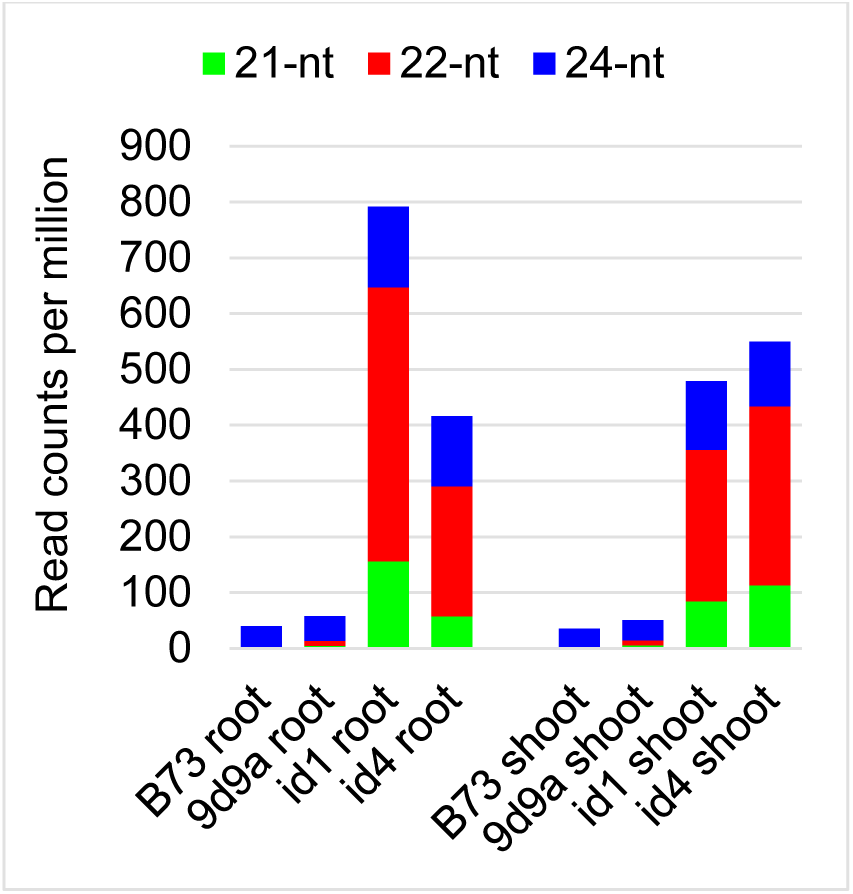
Small RNAs mapped to *Ac.* Compared to inbred B73 and progenitor allele *p1-vv-9D9A*, small RNA levels are increased significantly in both *p1-ww-id1* and *p1-ww-id4*. Colored bars indicate the abundance of 21-, 22- and 24-nt small RNA classes.

**Figure 8.**
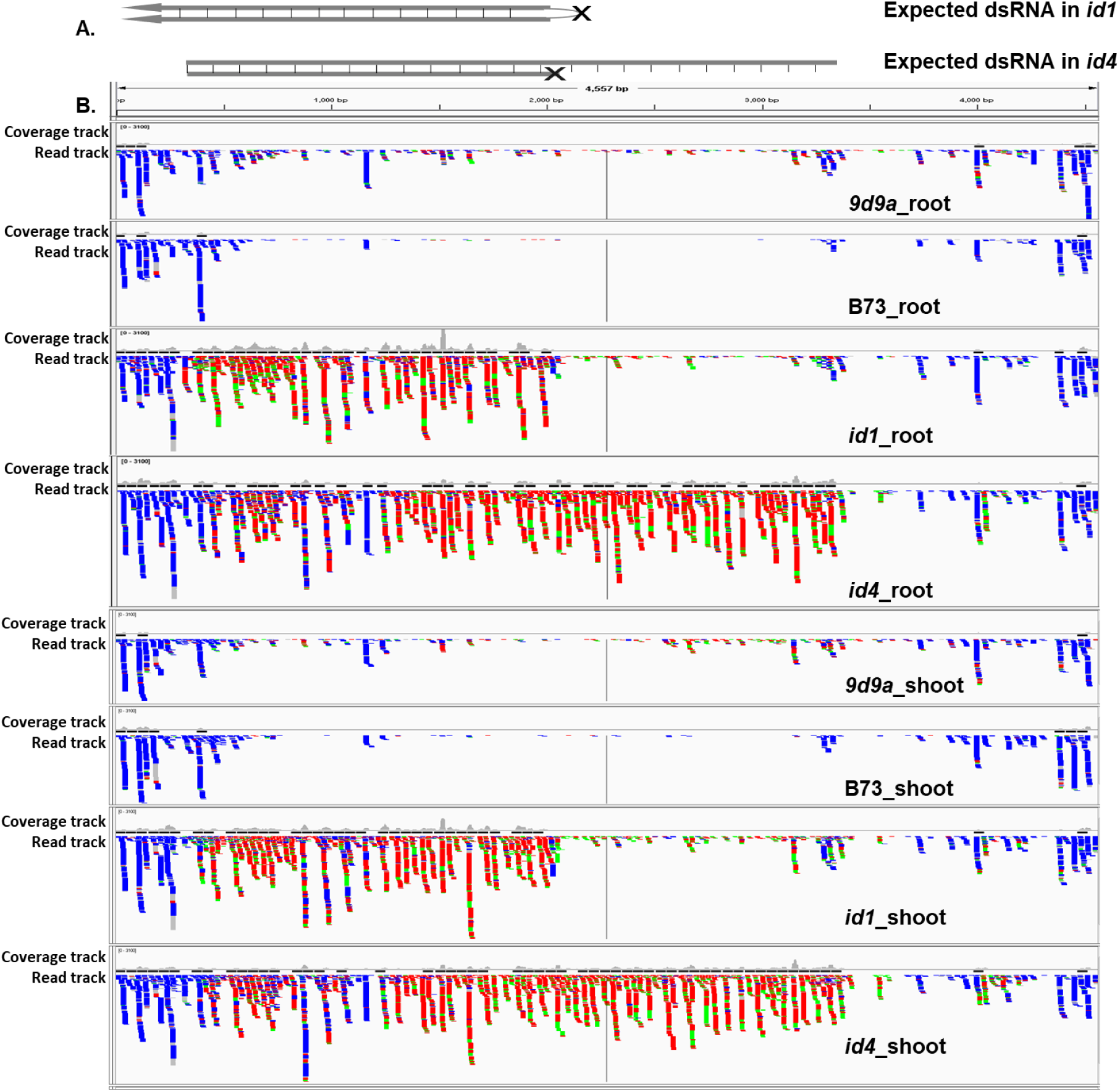
Detection and mapping of small RNA in *p1-ww-id1* and *p1-ww-id4*. **A.** The expected structures of dsRNA in *p1-ww-id1* and *p1-ww-id4*. The two inverted *fAc* fragments in *p1-ww-id1* are relatively symmetrical (2100-2486 nt of each indicated by Southern blot), predicting a hairpin structure in the dsRNA. The inverted *fAc* fragments in *p1-ww-id4* differ in size, with one *fAc* fragment of 1842-2486 nt and the other *fAc* of 2486-3900 nt indicated by Southern blot. This asymmetrical structure should give rise to a transcript with a stem and loop configuration in *p1-ww-id4*. **B.** Small RNAs mapped to *Ac* are visualized by the Integrative Genomics Viewer (Robinson et al. 2011). The y axis of the coverage track was standardized to the same scale for each sample. Colors indicate size of sRNA: Green, 21 nt; Red, 22 nt; Blue, 24 nt. In *p1-ww-id1* root, the enriched small RNAs map to a region 1-2100 nt from the *Ac* 5’ terminus, consistent with the dsRNA stem structure predicted by Southern blot. Small RNAs of the same profile were also observed in the shoot of *p1-ww-id1,* suggesting that small RNA may be transported to the shoot from the site of dsRNA synthesis in the root. In *p1-ww-id4,* the enriched small RNAs mapped to 1-3400 nt from the *Ac* 5’ terminus, consistent with the size of dsRNA predicted by Southern blots (2486 - 3900 bp) and indicating that small RNA is synthesized from both the double-strand stem and single-strand loop regions.

We observed similar profiles of siRNA in both root and shoot tissues of *p1-ww-id1,* even though no CI transcripts or dsRNAs were detected in *p1-ww-id1* shoots. Possibly, this result may reflect long-range transportation of siRNAs from root (where dsRNAs are produced) to shoot (where dsRNAs are absent) as earlier described in Arabidopsis (Chitwood and Timmermans 2010; Molnar *et al*. 2010). The *p1-ww-id4* dsRNAs are present in both shoot and root, thus we cannot distinguish whether the enriched siRNAs are from long-range transportation or local induction. In any event, the fact that the siRNAs correspond precisely to the structure of the two CIs strongly suggests that these CIs are the source of the siRNAs, and the production of siRNAs from the loop in the *p1-ww-id4* CI is consistent with the hypotheses that initial processing of the hairpin leads to small RNAs that retain the potential to trigger silencing of active elements *in trans*.

## Discussion

In this study, we characterized two naturally-occurring maize alleles (*p1-ww-id1* and *p1-ww-id4*) derived by Alternative Transposition (AT) of the transposable element *Ac.* Both alleles elicit similar *trans*-dominant and heritable repression of *Ac*-induced transposition and reduced levels of *Ac* mRNA. Both alleles contain inverted duplications and novel Composite Insertions (CIs) derived from SCT-induced DNA re-replication. The inverted duplications are of 3.2 Mb in *p1-ww-id1* and 0.3 Mb in *p1-ww-id4*; both CIs contain inverted repeats of the *Ac* 5’ terminal sequences, of 2.1-2.5 kb in *p1-ww-id1* and 1.8-3.9 kb in *p1-ww-id4*. These *Ac* inverted repeats are transcribed either from a flanking gene promoter in the case of *p1-ww-id1,* or likely from the *Ac* promoter itself in the case of *p1-ww-id4*. Double-stranded RNA transcripts were detected from both alleles, presumably due to fold back pairing of the inverted repeat transcripts.

The AT-initiated *de novo* and heritable silencing in the *p1-ww-id* alleles is consistent with mechanisms of TGS and PTGS reported for Class I elements in *Arabidopsis* (*de novo* silencing: Marí-Ordóñez et al. 2013b; Nuthikattu et al. 2013; McCue et al. 2014; Duan et al. 2015; Panda et al. 2016a; heritable silencing: Law and Jacobsen 2010; Haag and Pikaard 2011; Castel and Martienssen 2013; Matzke and Mosher 2014). In the *p1-ww-id* alleles described here, the 21-22 nt siRNAs enriched in the transcribed region of the CIs, and particularly in the loop region of *p1-ww-id4*, suggests the involvement of RDR6 in the *de novo* silencing. The reduced levels of *Ac* transcript can be formally explained by mRNA degradation via PTGS, and/or TGS of the *Ac* promoter triggered by the Pol II-RDR6 pathway. Of these mechanisms, TGS of *Ac* elements *in trans* seems most likely, as we see the heritable silencing of active *Ac* elements even after the *id1* and *id4* alleles are removed by meiotic segregation (Figure 2C-D). The production of 24-nt siRNAs corresponding to the TIR and sub-terminal regions of *Ac* signals the possible involvement of the Pol IV-RDR2 RdDM pathway in the maintenance of TGS by cytosine methylation in all sequence contexts (Law and Jacobsen 2010; Zemach *et al*. 2013; Stroud *et al*. 2014; Li *et al*. 2015). The siRNA-independent pathway may also participate in the maintenance of symmetrical methylation (Stroud *et al*. 2013).

The structure and effects of the *p1-ww-id1* and *p1-ww-id4* CIs are reminiscent of the *Mu killer* (*Muk*) allele that represses *Mutator* transposons in maize (Slotkin *et al*. 2003, 2005a; Li *et al*. 2010a), and hence the *p1-ww-id1* and *p1-ww-id4* alleles can be considered as examples of “*killers”* targeting a different DNA TE superfamily. *Mu killer* and the “*Ac killers*” share similar features: 1) they are both initiated from naturally occurred inverted duplications of partial DNA transposon sequences; 2) the transcription of inverted duplications in *Mu killer* and *Ac killer* in *p1-ww-id1* are both driven by nearby promoters, while *p1-ww-id4* is apparently transcribed by the *Ac* promoter, producing dsRNAs as the precursor of 21, 22, and 24-nt siRNAs; 3) both can trigger *in-trans* and heritable silencing of an active element; and 4) there is evidence for long-range transportation of siRNAs in both systems. These similarities suggest a general mechanism for the heritable silencing of active DNA transposons.

Our findings of siRNA-mediated *de novo* and heritable silencing of *Ac/Ds* adds another layer of complexity to the regulation of *Ac/Ds* activity. Previous studies of the regulation of *Ac/Ds* activity reveal a complex relationship among *Ac* dosage, transcript abundance, transposase level, DNA methylation and transposition frequency. *Ac* has a GC-rich subterminal region, and hypermethylation of this region is associated with transcriptional silencing (Kunze *et al*. 1988; Brutnell and Dellaporta 1994; Conrad and Brutnell 2005). Additionally, splicing of *Ac* mRNA is inefficient and inaccurate, producing a large number of aberrant *Ac* transcripts in *Arabidopsis*, representing post-transcriptional regulation (Jarvis *et al*. 1997). The aggregation of *Ac* transposase protein in tobacco (Kunze *et al*. 1995), petunia and maize (Heinlem *et al*. 1994), may represent a type of post-translational regulation responsible for the *Ac* negative dosage effect first observed by McClintock (McClintock 1948, 1949, 1950). Molecular analysis showed that increased dosage of *Ac* results in increased *Ac* transposase mRNA and protein levels, at least in the maize *wx-m7* allele that was tested (Fußwinkel et al., 1991). In this study, we find that the *p1-ww-id1* and *p1-ww-id4* alleles derived from AT-induced DNA re-replication unveil yet another level of complexity, involving the transgenerational silencing of active elements via *trans*-acting siRNAs.

This study also expands our understanding of the impact of AT on the structure of the maize genome. Previous studies have described the ability of AT to induce structural rearrangements such as deletions, inversions, translocations, duplications (Zhang and Peterson 2004; Huang and Dooner 2008; Zhang *et al*. 2009, 2013) and exon shuffling (Zhang et al. 2006). Here we show that *p1-ww-id* alleles derived from SCT contain novel inverted-repeat structures (CIs) generated by DNA re-replication. The fact that similar structures and mechanisms effect silencing in members of two unrelated transposon superfamilies (*hAT* and *Mu*) suggests that IR-induced siRNA may represent a general mechanism of spontaneous TE silencing.

## Author Contributions

DW, JZ and TP conceived and designed the experiment; DW performed the experiments; TZ, MZ and DW analyzed the small RNAseq data; DW, DL, and TP wrote the paper.

## Acknowledgements

This research is supported by National Science Foundation award 0923826 (to T.P. and J.Z.), by the USDA National Institute of Food and Agriculture Hatch project number IOW05282, and by State of Iowa funds. We thank Dr. R. Keith Slotkin for suggestions on dsRNA experiment. We thank Lisa Coffey and Dr. Patrick Schnable for providing growth chamber. We thank Terry Olson for technical assistance, and Douglas Baker for field assistance.

## Video 1

## Supplemental Material

### a. Figures

**Figure S1.**
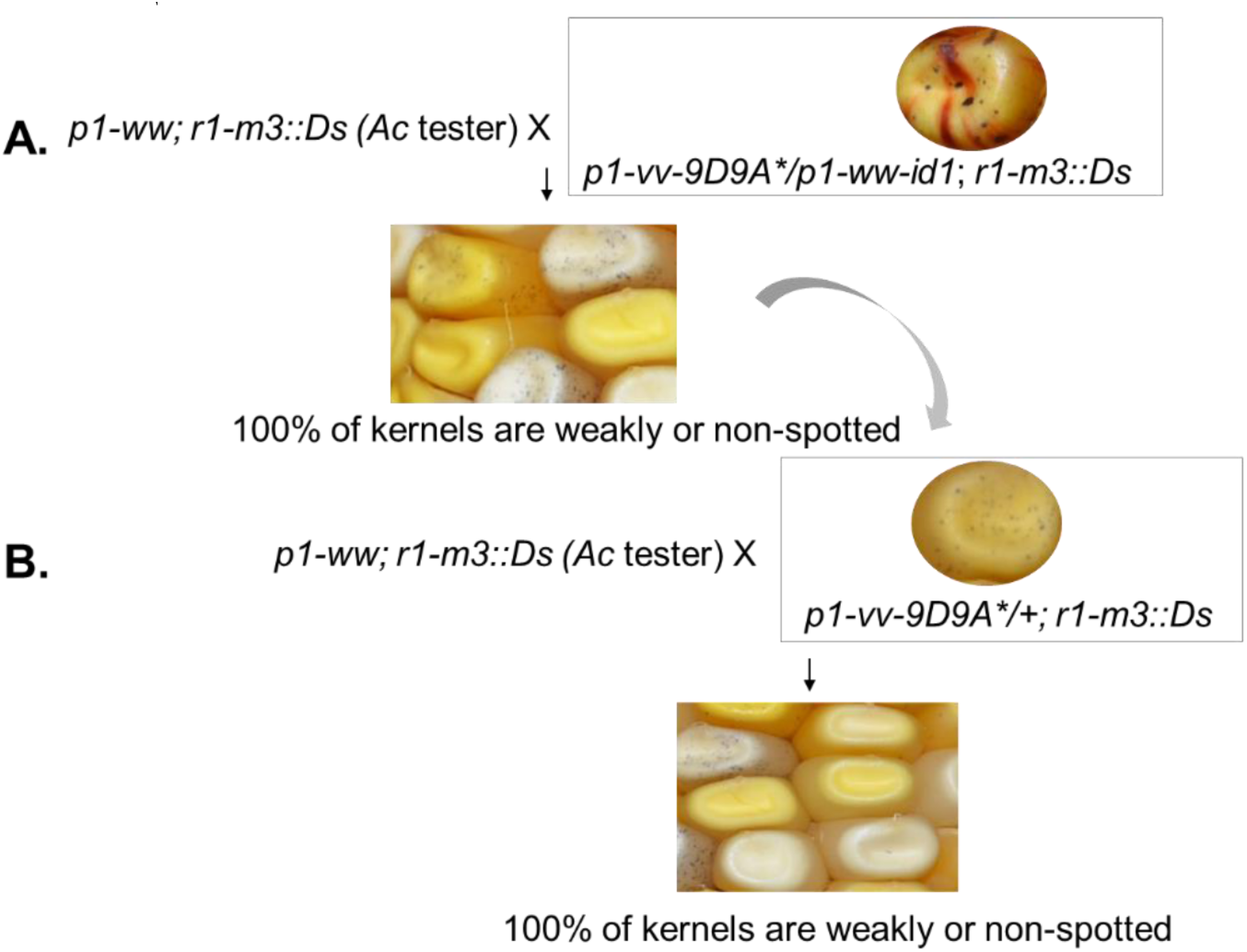
Genetic crosses to test the maintenance of *Ac* repression initiated by *p1-ww-id1*. **A.** The ear from the cross of *Ac* tester by *p1-vv-9D9A*/p1-ww-id1.* 100% of the kernels show fine spots or no spots on the aleurone, indicating that the silencing of *Ac* by *p1-ww-id1* was efficiently maintained in the first-generation following segregation of *p1-ww-id1* from *p1-vv-9D9A*.* Asterisk indicates silenced *Ac*. **B.** The ear from the cross of *Ac* tester by *p1-vv-9D9A*/+.* 100% of the kernels show fine spots or no spots on the aleurone, indicating that the silencing of *Ac* by *p1-ww-id1* was efficiently maintained for the second generation in the absence of *p1-ww-id1*.

**Figure S2.**
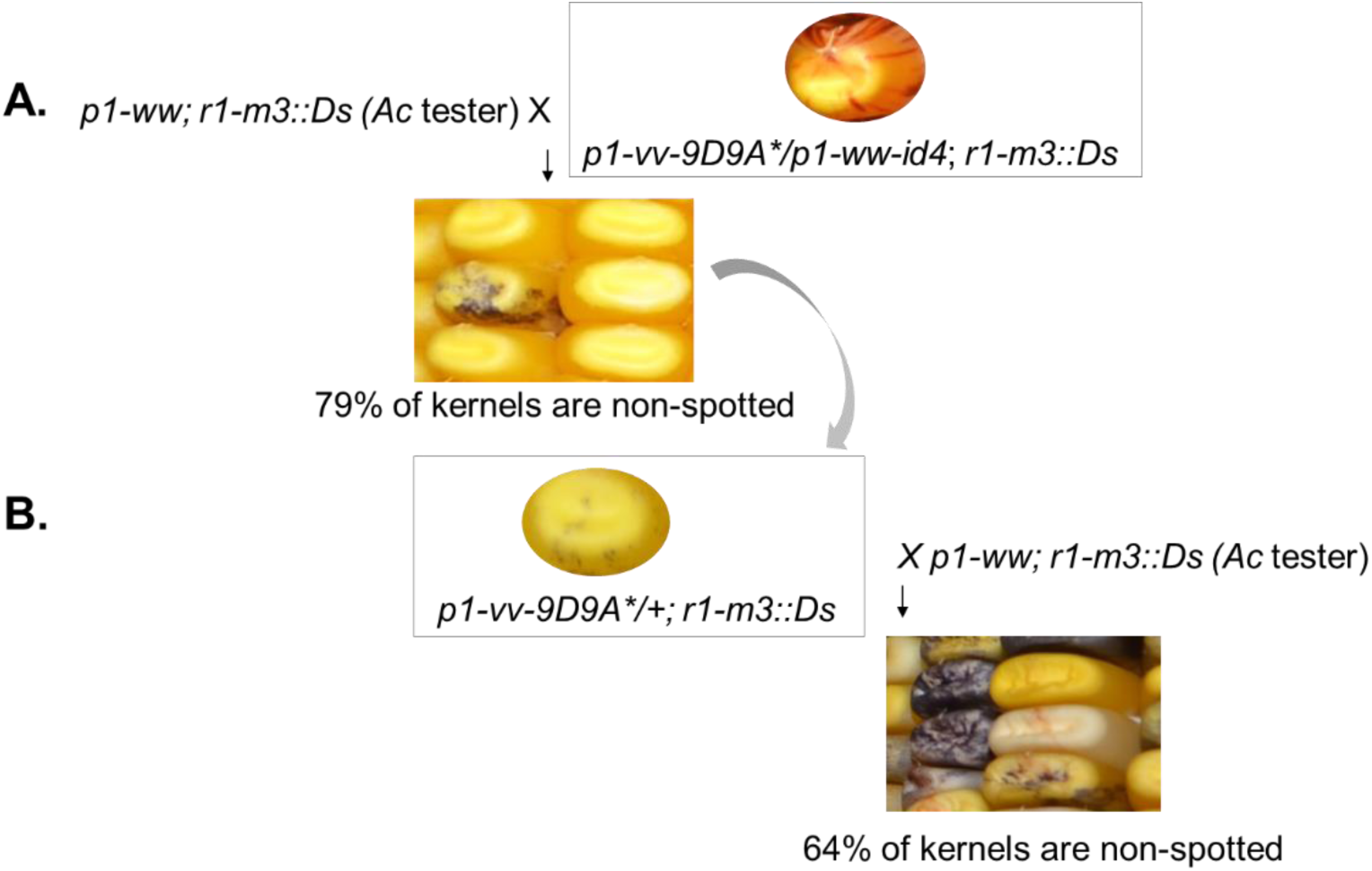
Genetic crosses to test the maintenance of *Ac* repression initiated by *p1-ww-id4*. **A.** The ear from the cross of *Ac* tester by *p1-vv-9D9A*/p1-ww-id4.* 79% of kernels show fine spots or no spots on the aleurone, indicating that silencing was maintained on 58% of *Ac* elements exposed to *p1-ww-id4.* Asterisk indicates silenced *Ac*. **B.** The ear from the cross of *Ac* tester by *p1-vv-9D9A*/+.* 64% of kernels show fine spots or no spots on the aleurone, indicating that silencing was maintained on 28% of *Ac* elements in the second-generation following exposure to *p1-ww-id4*.

**Figure S3.**
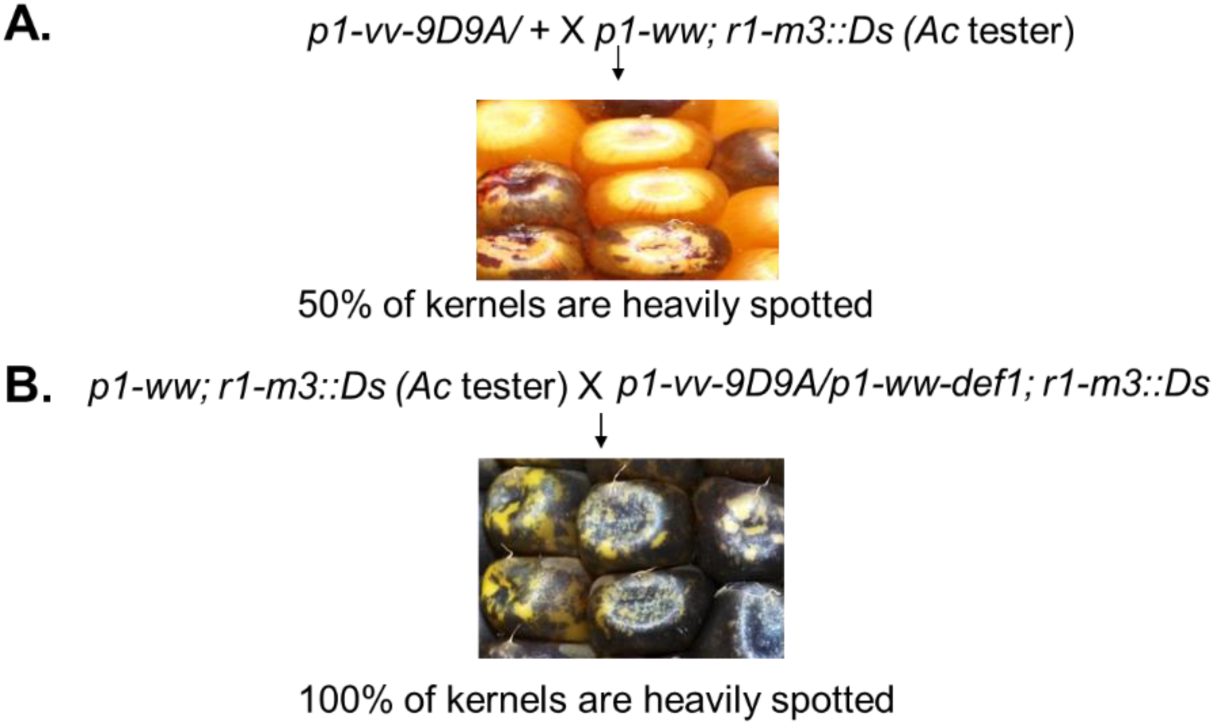
Genetic crosses to test the activity of *Ac* in the progenitor *p1-vv-9D9A*. **A.** The ear from the cross of *p1-vv-9D9A*/+ by *Ac* tester (*p1-ww; r1-m3::Ds*). About 50% of the kernels show heavy spotting on the aleurone, consistent with the segregation of one active *Ac* element from the heterozygous progenitor *p1-vv-9D9A/+*. **B.** The ear from the cross of *p1-vv-9D9A*/*p1-ww-def1* by *Ac* tester (*p1-ww; r1-m3::Ds*). All the kernels show heavy spotting on the aleurone, consistent with segregation of two active *Ac* elements at allelic positions (one each at *p1-vv-9D9A* and *p1-ww-def1*).

**Figure S4.**
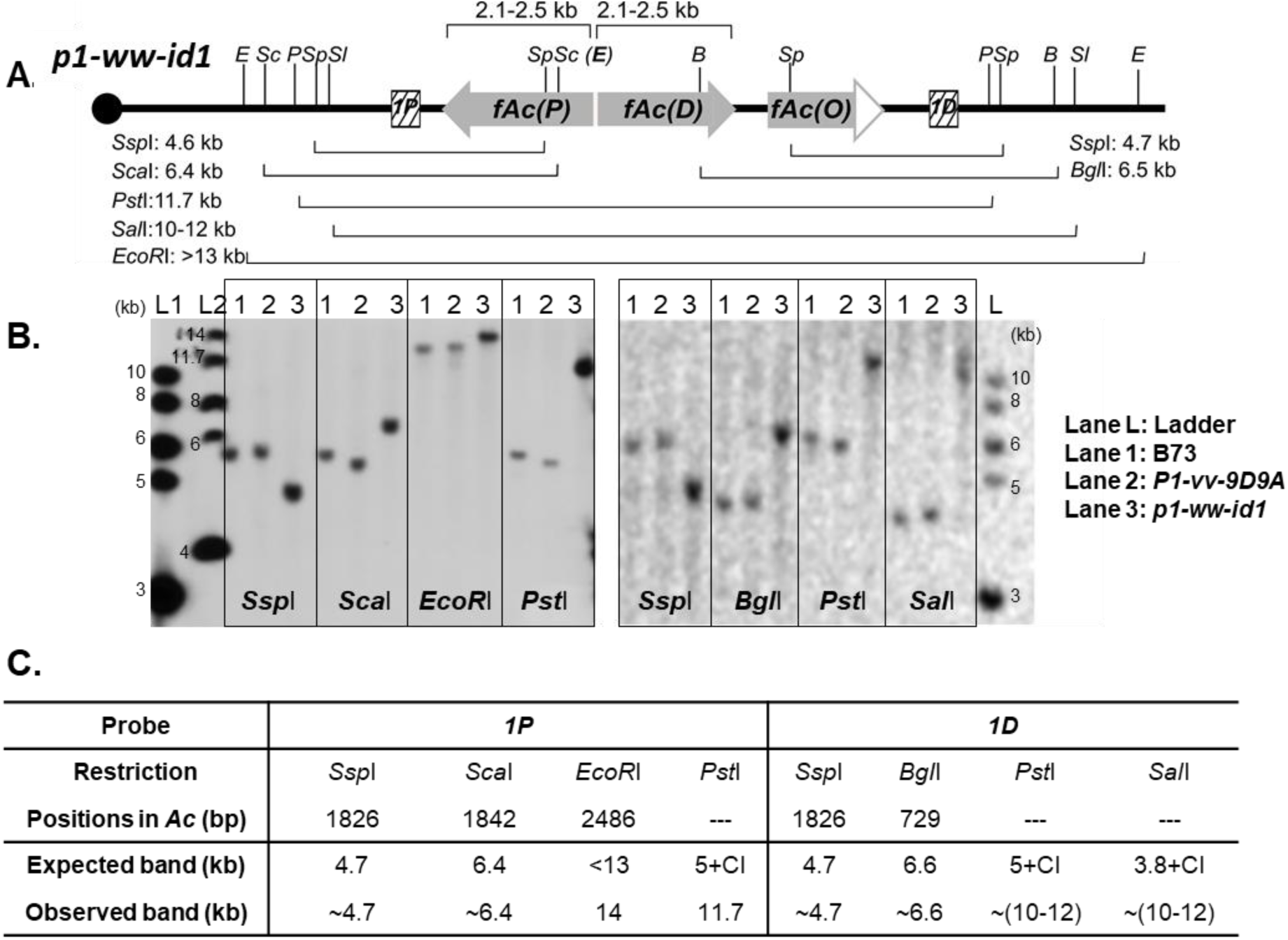
Southern blot analysis of the structure of Composite Insertion of *p1-ww-id1*. **A.** Schematic structure of the *p1-ww-id1* allele with restriction sites (vertical lines) and probes (hatched boxes marked “*1P*” and “*1D*”) indicated. **B.** The Southern blot result of genomic DNAs from B73 (Lane 1), *p1-vv-9D9A* (Lane 2), and *p1-ww-id1* (Lane 3), digested with the indicated enzymes and hybridized with probe *1P* (left panel) and *1D* (right panel). **C.** Table of expected and observed band sizes when the restriction sites are included in the CI. Enzyme Abbreviation: E: *EcoR*I; Sc: *Sca*I; P:*Pst*I; Sp: *Ssp*I; Sl: *Sal*I; B: *Bgl*I; N: *Nco*I.

**Figure S5.**
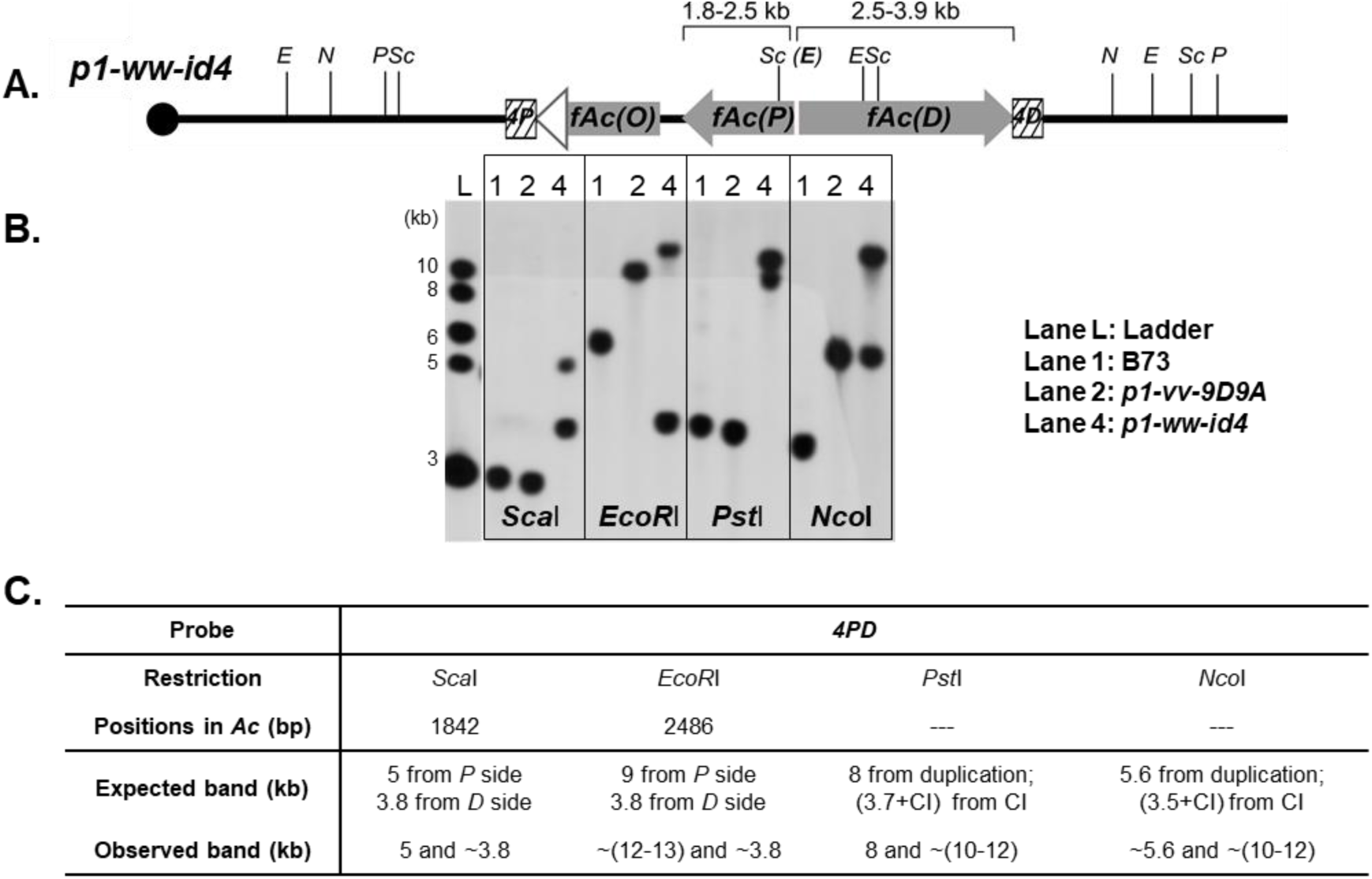
Southern blot analysis of the structure of Composite Insertion in *p1-ww-id4*. **A.** Schematic structure of the *p1-ww-id4* allele with restriction sites (vertical lines) and probes (hatched boxes labeled “*4P*” and “*4D*”) indicated. Enzyme Abbreviation: E: *EcoR*I; N: *Nco*I; P: *Pst*I; Sc: *Sca*I. **B.** Southern blot result of genomic DNA from B73 (Lane 1), *p1-vv-9D9A* (Lane 2), and *p1-ww-id4* (Lane 4), digested with the indicated enzymes and hybridized with probe *4PD*. **C.** Table of expected and observed band sizes when the restriction sites are included in the CI. Enzyme Abbreviation: E: *EcoR*I; Sc: *Sca*I; P:*Pst*I; Sp: *Ssp*I; Sl: *Sal*I; B: *Bgl*I; N: *Nco*I.

**Figure S6.**
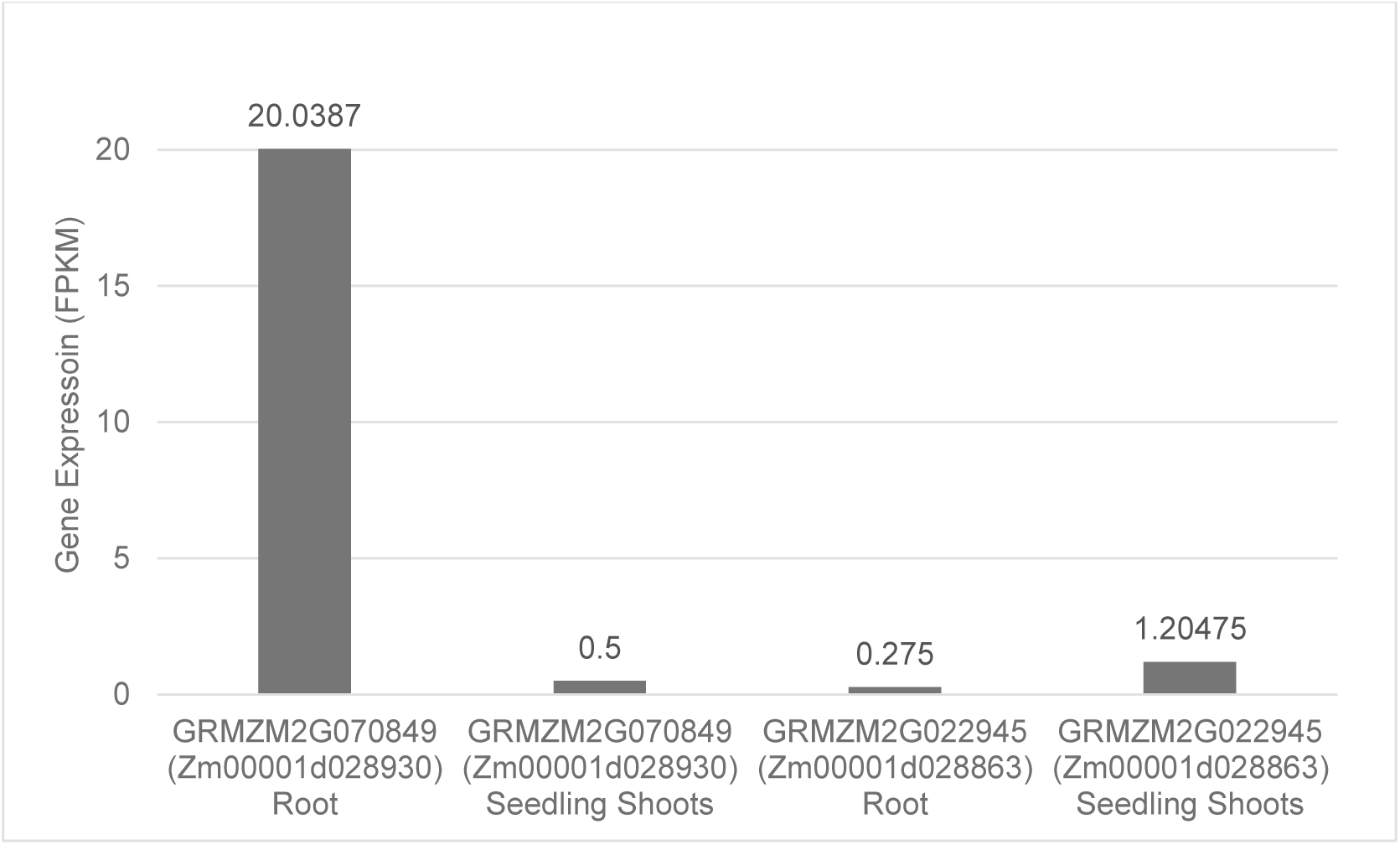
The gene expression data of target genes where CI-*id1* and CI-*id4* are inserted. Data was extracted from Wang 2009 (Wang et al. 2009) paper.

**Figure S7.**
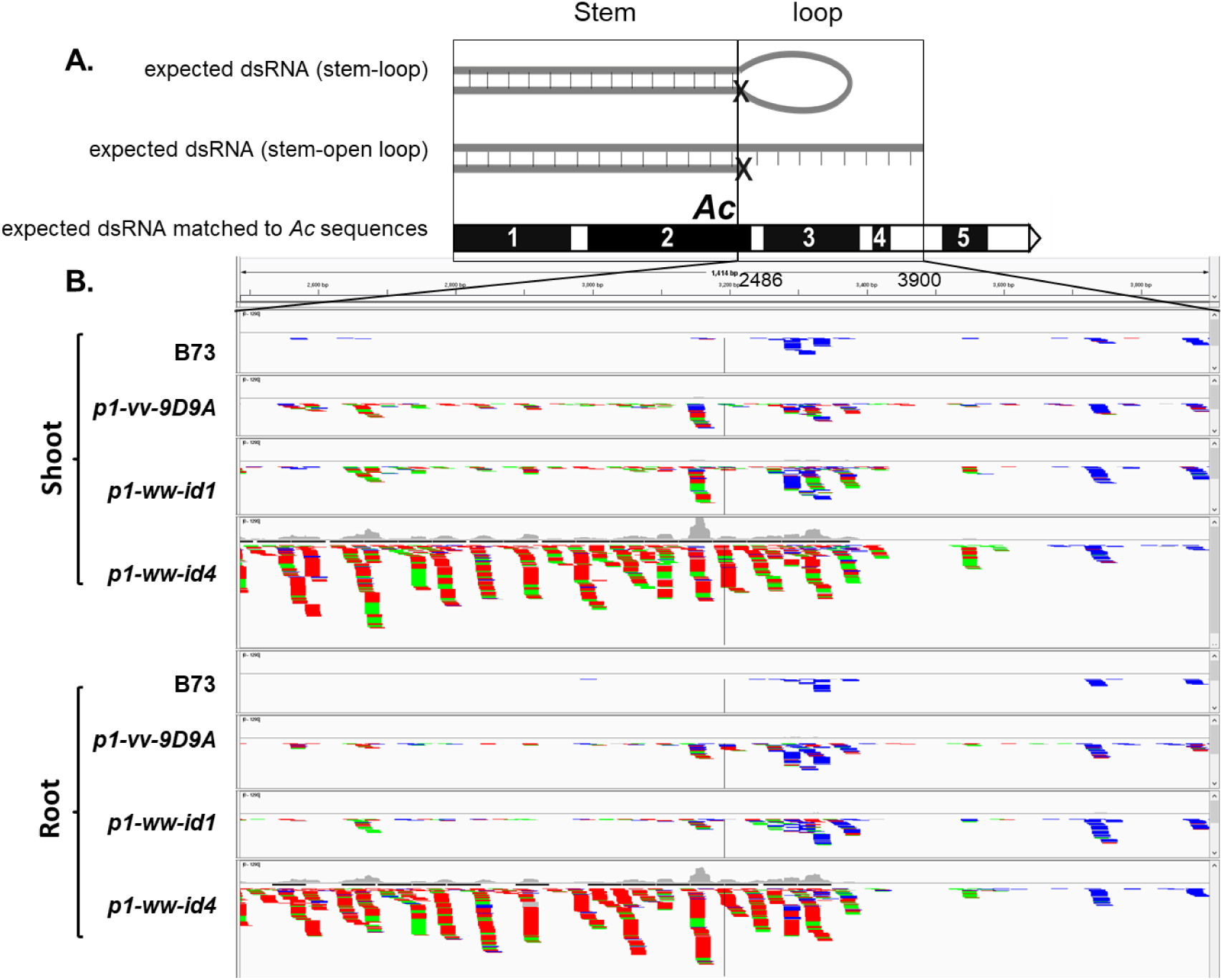
Small RNAs mapped to the loop region of *p1-ww-id4*, visualized by the Integrative Genomics Viewer. **A.** Diagram of expected dsRNA from *p1-ww-id4* including predicted stem-loop structure (top), and stem-open loop structure to show extent of *Ac* sequence included in loop (middle). Lower map shows structure of *Ac* including exons 1 – 5 (black boxes), and predicted loop positions at 2486 – 3900. **B.** Small RNA reads mapped to *Ac* positions 2486 – 3900 in shoot and root tissues of the indicated genotypes. Green, red and blue indicate 21-, 22-, and 24-nt small RNAs, respectively. Compared to genotypes that do not contain a loop structure (B73, progenitor allele *p1-vv-9D9A* and *p1-ww-id1*), the *p1-ww-id4* allele shows an accumulation of 21-22 nt small RNAs in both shoot and root. The small RNAs in *p1-ww-id4* extend from 2486 to ∼3400 in *Ac*, which is similar to the size of loop predicted by genomic Southern blot (2486 – 3900). The ∼500 bp difference may due to the low resolution of genomic Southern blot in determining fragment sizes.

**Figure S8.**
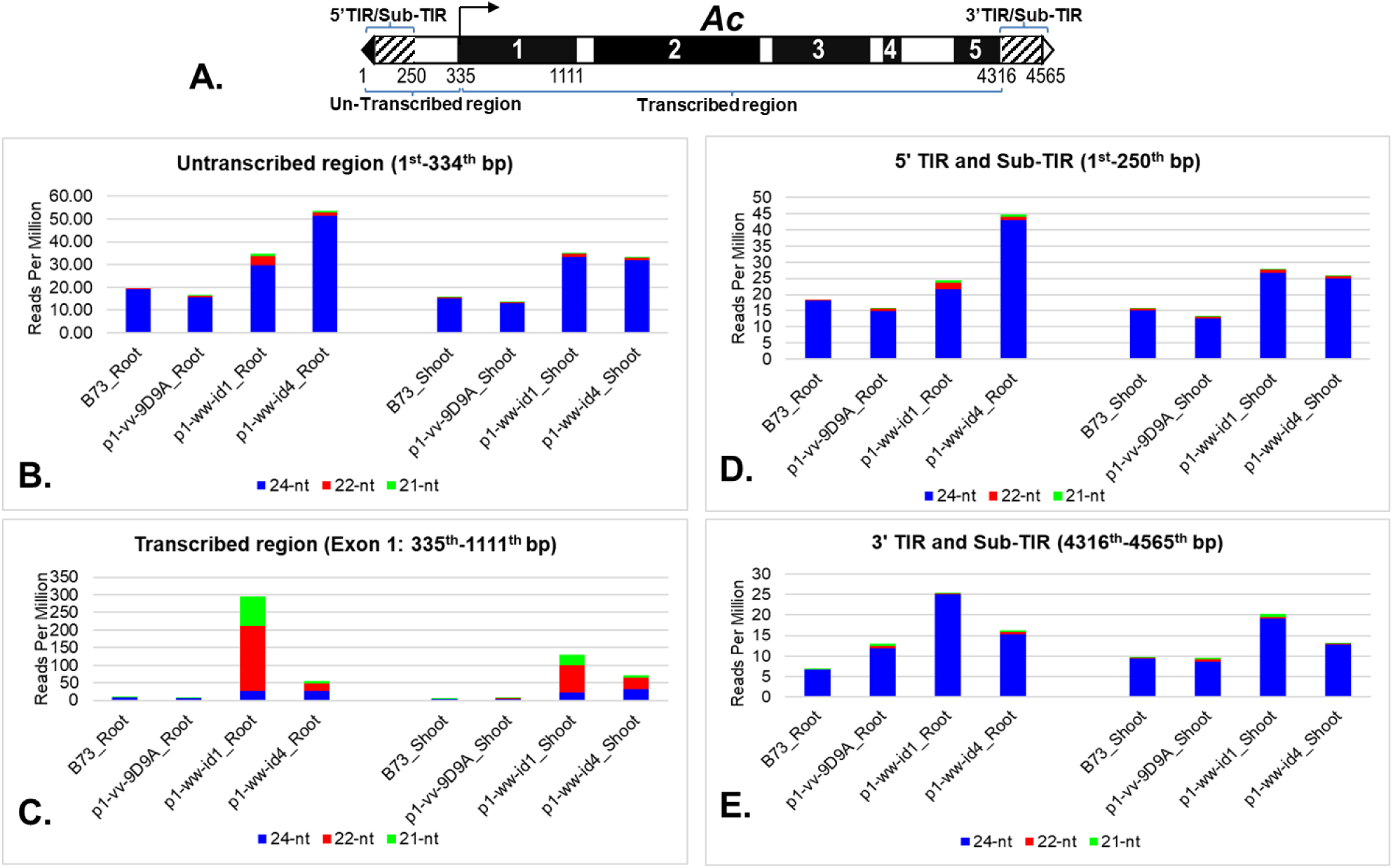
Small RNA read counts of specific regions of *Ac*. **A.** Schematic structure of *Ac.* Filled and open triangles indicate *Ac* 5’ and 3’ termini, respectively. Filled boxes are exons, hatched boxes indicate sub-terminal regions. The arrow marks the transcription start site of *Ac*, and numbers indicate the nucleotide positions within *Ac*, including untranscribed region (1-334 bp; 5’ TIR/Sub-TIR from 1 – 250 bp); transcribed region (335 – 4315; exon 1 from 335 – 1111 bp); 3’ TIR/Sub-TIR (4316 – 4565 bp). **B-E.** Small RNA Reads Per Million that exactly match the *Ac* sequence of the indicated regions of *Ac*.

**Figure S9.**
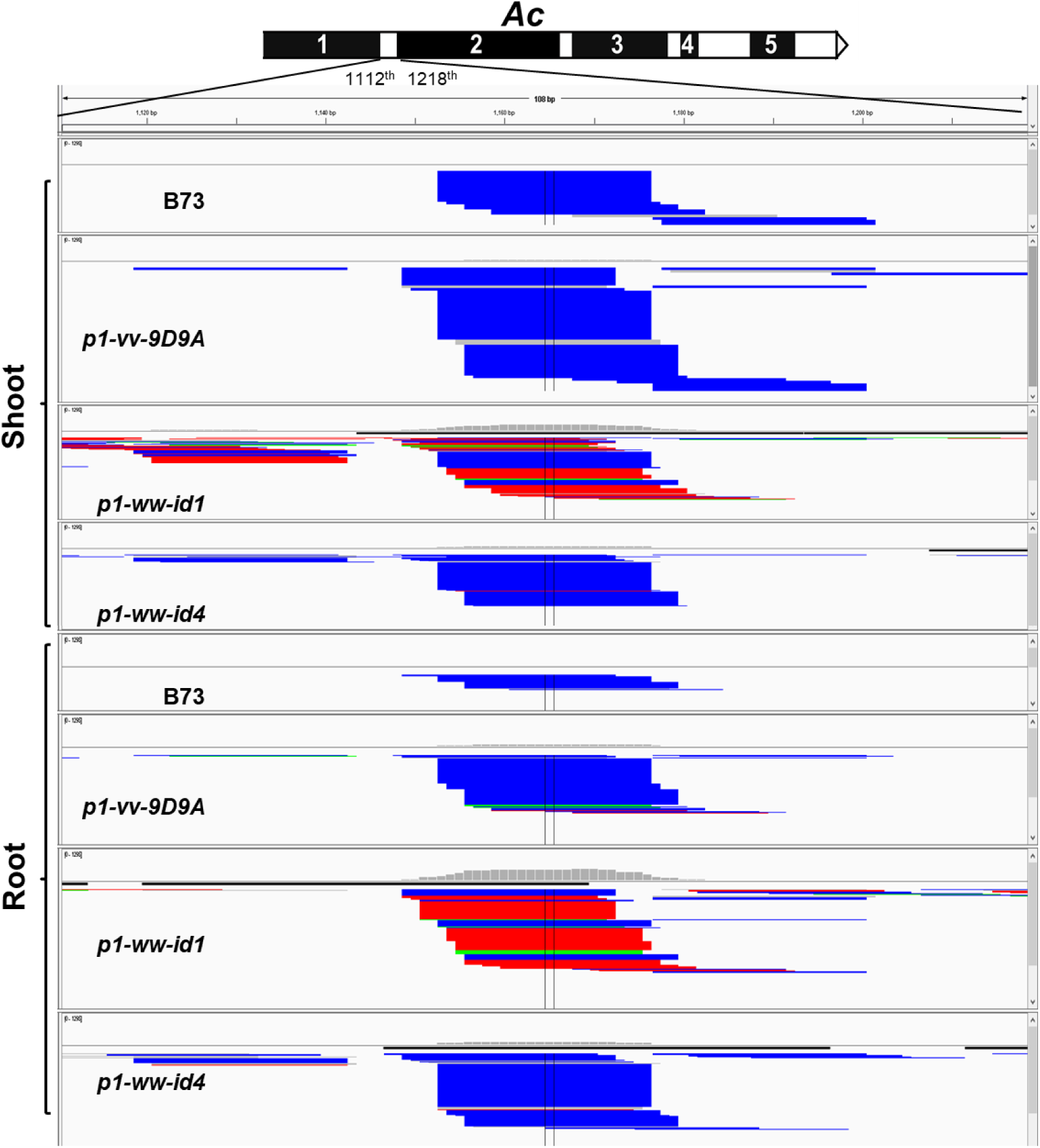
Small RNAs mapped to *Ac* intron 1 (position 1112 -1218 bp), visualized by the Integrative Genomics Viewer. The y axis of the coverage track was standardized to the same scale for each sample. Green, red and blue indicate 21-, 22-, and 24-nt small RNAs, respectively. The 21- and 22-nt small RNAs are enriched in both root and shoot of *p1-ww-id1,* consistent with the intron-containing cDNA that was detected by RT-PCR. The extent of small RNA coverage also matches the size of unspliced intron 1 in *p1-ww-id1*. Unexpectedly, *p1-ww-id4* does not show a similar accumulation of 21- and 22-nt sRNA, even though it does produce dsRNA including the unspliced *Ac* intron 1 based on RT-PCR and dsRNA protections assays (Figure 5). The reason for this difference is unknown.

### b. Tables

**Table S1.** Kernel counts from the cobs of the indicated crosses. *p1-vv-9D9A\** was segregated from *p1-vv-9D9A/p1-ww-id*. ^†^In this cross, *p1-vv-9D9A\** was segregated from *p1-vv-9D9A/p1-ww-id1*. **^§^** In this cross, *p1-vv-9D9A\** was segregated from *p1-vv-9D9A/p1-ww-id4*

| Crosses | coarsely spotted | fine spotted | no spots | total seeds |
| --- | --- | --- | --- | --- |
| *p1-vv-9D9A* x *p1-ww; r1-m3::Ds* (Ac tester) | 112 | 0 | 121 | 233 |
| *p1-ww; r1-m3::Ds* (Ac tester) x *p1-ww-id1* | 4 | 38 | 97 | 139 |
| *p1-ww; r1-m3::Ds* (Ac tester) x *p1-ww-id4* | 0 | 0 | 21 | 21 |
| *p1-vv-9D9A* x *p1-ww-id1; r1-m3::Ds* | 11 | 5 | 82 | 98 |
| *p1-vv-9D9A* x *p1-ww-id4; r1-m3::Ds* | 4 | 0 | 18 | 22 |
| *p1-ww; r1-m3::Ds* (Ac tester) x *p1-vv-9D9A/p1-ww-def1; r1-m3::Ds* | 112 | 0 | 0 | 112 |
| *p1-ww; r1-m3::Ds* (Ac tester) x *p1-vv-9D9A/p1-ww-id1; r1-m3::Ds* | 0 | 16 | 24 | 40 |
| *p1-ww; r1-m3::Ds* (Ac tester) x *p1-vv-9D9A\**; *r1-m3::Ds* † | 0 | 64 | 224 | 288 |
| *p1-ww; r1-m3::Ds* (Ac tester) x *p1-vv-9D9A/p1-ww-id4; r1-m3::Ds* | 34 | 21 | 105 | 160 |
| *p1-ww; r1-m3::Ds* (Ac tester) x *p1-vv-9D9A\**; *r1-m3::Ds* § | 61 | 5 | 104 | 170 |

**Table S2.**
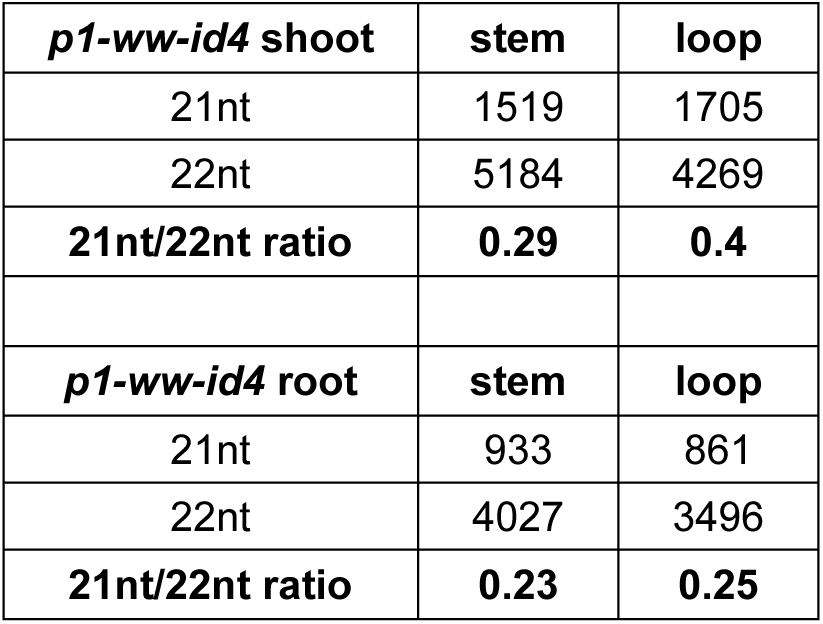
21 and 22nt small RNA read counts in the stem and loop region of dsRNA in *p1-ww-id4*. Chi-squared test p < 0.01 in shoot and p =0.24 in root.

**Table S3.**
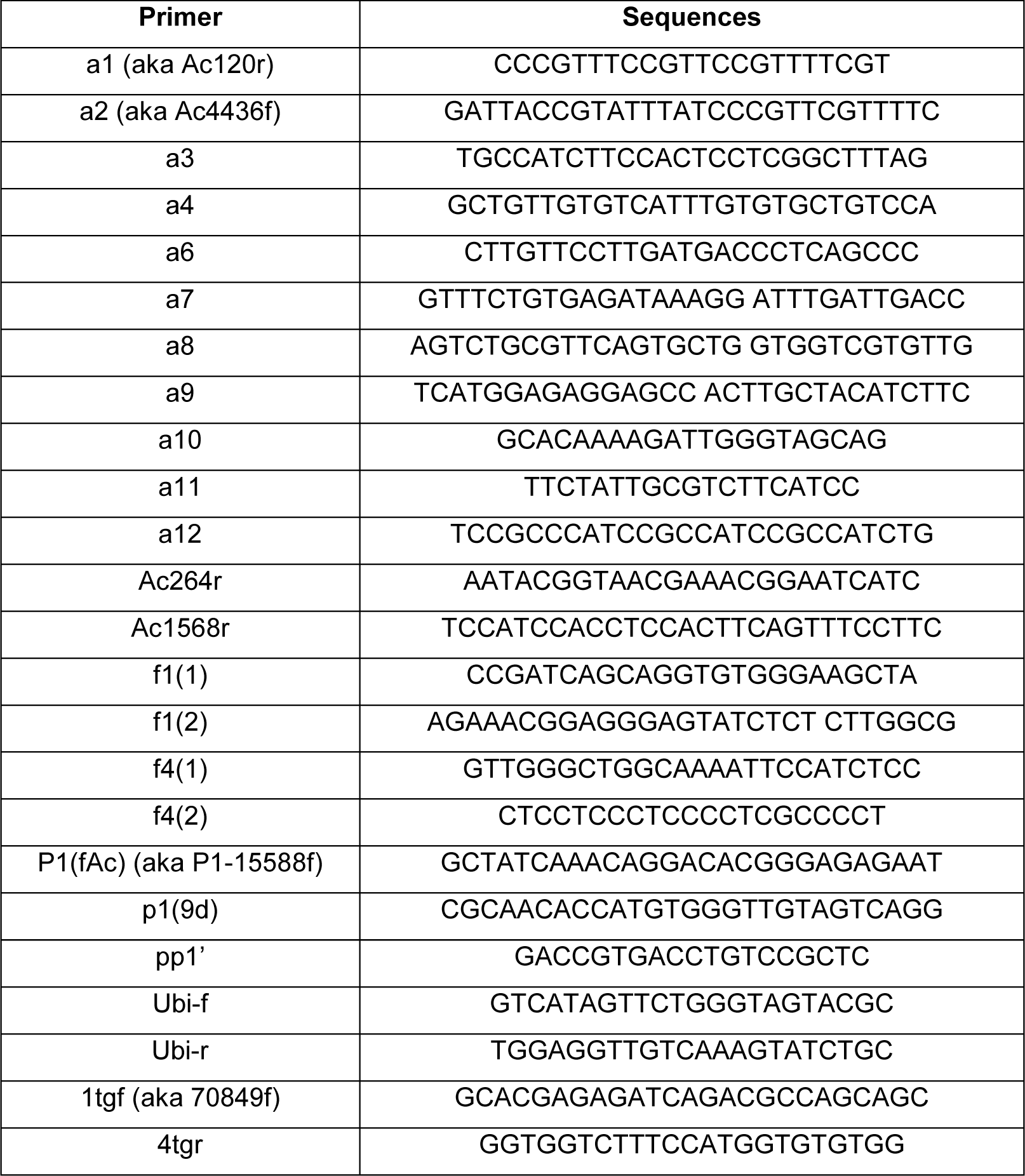
Oligonucleotide primer sequences used in PCR experiments.

### c. Files

**File S1.** Genomic sequences of *p1-ww-id1* and *p1-ww-id4* at site of excision of *Ac* and *fAc* termini. Fragments were obtained from genomic PCR and sequenced using indicated primer *P1-*15588F. The 2 bp “footprints” remaining after SCT excision are shown in red font. Sequences of the progenitor *p1-vv-9D9A* allele are shown for comparison.

> *p1-ww-id1*_*P1*-15588F

NNNNNNNNNANNNATCCNCGTGCATGCATGCCACTGTAGCGCCGTAATATAATGATAGATA TGCGCTATTGCTCCTACAACTACAAGTCTACAACCCACATGGTGTTGCGAGAGCTAGCGGTG CCACACAGTCATGGAAACATGGTTTGTGAAAGCAGCTTAACTAATTACTAGCTTAACTAATT ACTAACTAATTACTAGCCTGCACTAATAAGGCTTAAAACAAGTGATCCTCGCAGGTATGTTT GTCTCAATTGTTGTACATGTCATCATTATAAATTCTCAATTAATCAAATGTCAATTATTGTAG GTACGATGCAATTTGTCCTAAAGCCGAGGAGTGGAAAGATGGCAA

> *p1-ww-id4*_*P1*-15588F

NNNNNNNNNNNNNNNTCCNCGTGCATGCATGCCACTGTAGCGCCGTAATATAATGATAGAT ATGCGCTATTGCTCCTACAACTACAAGTCTACAACCCACATGGTGTTGCGAGAGCTAGCGGT GCCACACAGTCATGGAAACATGGTTTGTGAAAGCAGCTTAACTAATTACTAGCTTAACTAAT TACTAACTAATTACTAGCCTGCACTAATAAGGCTTAAAACAAGTGATCCTCGCAGGTATGTT TGTCTCAATTGTTGTACATGTCATCATTATAAATTCTCAATTAATCAAATGTCAATTATTGTA GGTACGATGCAATTTGTCCTAAAGCCGAGGAGTGGAAGATGGCAA

>*p1-vv-9D9A*

AGAAATCATCTAACAAAACTGGCGAGCTATCAAACAGGACACGGGAGAGAATAGATGATTA AACAATAATCCCTCGTGCAATGCATGCCACTGTAGCGCCGTAATATAATGATAGATATGCGC TATTGCTCCTACAACTACAA**CA**CTACAACCCACATGGTGTTGCGAGAGCTAGCGGTGCCACA CAGTCATGGAAACATGGTTTGTGAAAGCAGCTTAACTAATTACTAGCTTAACTAATTACTAA CTAATTACTAGCCTGCACTAATAAGGCTTAAAACAAGTGATCCTCGCAGGTATGTTTGTCTC AATTGTTGTACATGTCATCATTATAAATTCTCAATTAATCAAA<colcnt=3>

id1 CGCTATTGCTCCTACAACTACAAGTCTACAACCCACATGGTGTTGCGAGAGCTAGCGGTG 123

id4 CGCTATTGCTCCTACAACTACAAGTCTACAACCCACATGGTGTTGCGAGAGCTAGCGGTG 124

9D9A CGCTATTGCTCCTACAACTACAA**CA**CTACAACCCACATGGTGTTGCGAGAGCTAGCGGTG 180

*********************** ***********************************

**File S2.** Genomic sequences showing junctions of CI termini and flanking DNA in *p1-ww-id1* and *p1-ww-id4*. Fragments were obtained from genomic PCR and sequenced using indicated primers Ac264r, Ac4436f, and Ac120r. Underlined regions indicate *Ac* or *fAc;* motifs in yellow highlight indicate 8 bp TSDs flanking each CI insertion. Note that the 8 bp TSDs for each allele are shown as reverse complements due to the opposite orientation of sequencing reactions.

> *p1-ww-id1_*Ac_Ac264r_912121.seq

<u>NNNNNNNNNNGNTTCGTTCGTTTTCGTTTTTTACCTCGGGTTCGAAATCGATCGGGATAAAACTAACAAAATCGGTTATACGATAACGGTCGGTACGGGATTTTCCCATCCTACTTTCATCCCTG</u> AGCGAGGCGGCGGCTACGAATTGTGGCTGCTGTTAGTTGTTGCTCAACTCACGATGATGCAA AGCTAGCTAGCTAGCATCGATCGATCGGCATCATGCAATTTGATGACGATCCTAGCTAGCTC TAGCTACATGTATATGCATATCCTAGGCAGCTAGCTTCCCACACCNGGCTGATCGGN

> *p1-ww-id1_*fAc_Ac4436f_912122.seq

<u>NNNNNNNNNNNNNNTTTTCGTNNNNNNNNCAAGTTAAATATGAAAATGAAAACGGTAGAG</u> GTATTTTACCGACCGTTACCGACCGTTTTCATCCCTAGCCTCGCTCGCTCATGGGTTCTATCACATCGCGCGCCCACAGATGCTCCACAGCTTTACTTGCGCATATCGTTTTCTTCTCCGCGTTGCCGTACAAGCCTGGGCAATCGTGGTGGCACGTGCACGTTCATGCTAATATGGAAACAGATAGGCTAGCTGCTAGGTAGGAAACCATCCCAGCTAACCAGAGCCGCCAAGAGAGATACTCCCTCCGTTNNNNN

> *p1-ww-id4*_Ac120r_793173.seq

<u>NNNNNNNNNNNNNNCGNNCGNNNNNACTAACANAATCGGTTATACGATAACGGTCGGTAC</u> GGGATTTTCCCATCCTACTTTCATCCCTGGCCCGGATGCTGATGTAGCGACGATGCCCTGTAC GGAACGATGATGGTGACGGCACGCATCGGATTCCATTCCGTTCCAGCGGGAGGGGCGAGGG GAGGGAGGAG

> *p1-ww-id4*_ac4436f_792285.seq

<u>NNNNNNNNNNNNNNTNTTCGTNNNNNNNNNAAGTTAAATATGAAAATGAAAACGGTAGAG</u> GTATTTTANNNNNNNNNANNGACCGTTTTCATCCCTAATCCGGGCGCTGATGGGGGTGTCTC CCTAGTAGTTTTAAAATGCCAATGTAATCAGCGTCTTTTTAGATGCGAAAGCCAGCCAGCTC CCATCCAGTTGAAGCGATGACGACGGGCGCAAGAGAAAGAAAAGCATGTATGCGTGCCAAA TTAAAGCAAGGAGATGGAATTTTGCCAGCCCAACGAAAAGGCCATTGTGCGATTCAGCGGC TCATTAAATTATTGGCTCTCTGTTCTGTTCACATCTGAACGAACCACCAGCGTGGCGTACGAC CGTGATTCTTCTTGTCAAGACAGT

**File S3.** Sequences of *p1-ww-id1* root chimeric transcript.

**A.** CI transcript in *p1-ww-id1* includes Intron 1 of host gene Zm00001d028930. Sequences were obtained from 5’ region of *p1-ww-id1* root transcript using primer 70849f located in Exon 1 of host gene Zm00001d028930. id1-c: cDNA; id1-g: genomic DNA.

Exon of Zm00001d028930

Intron of Zm00001d028930

Ac TIR

*Ac* Intron

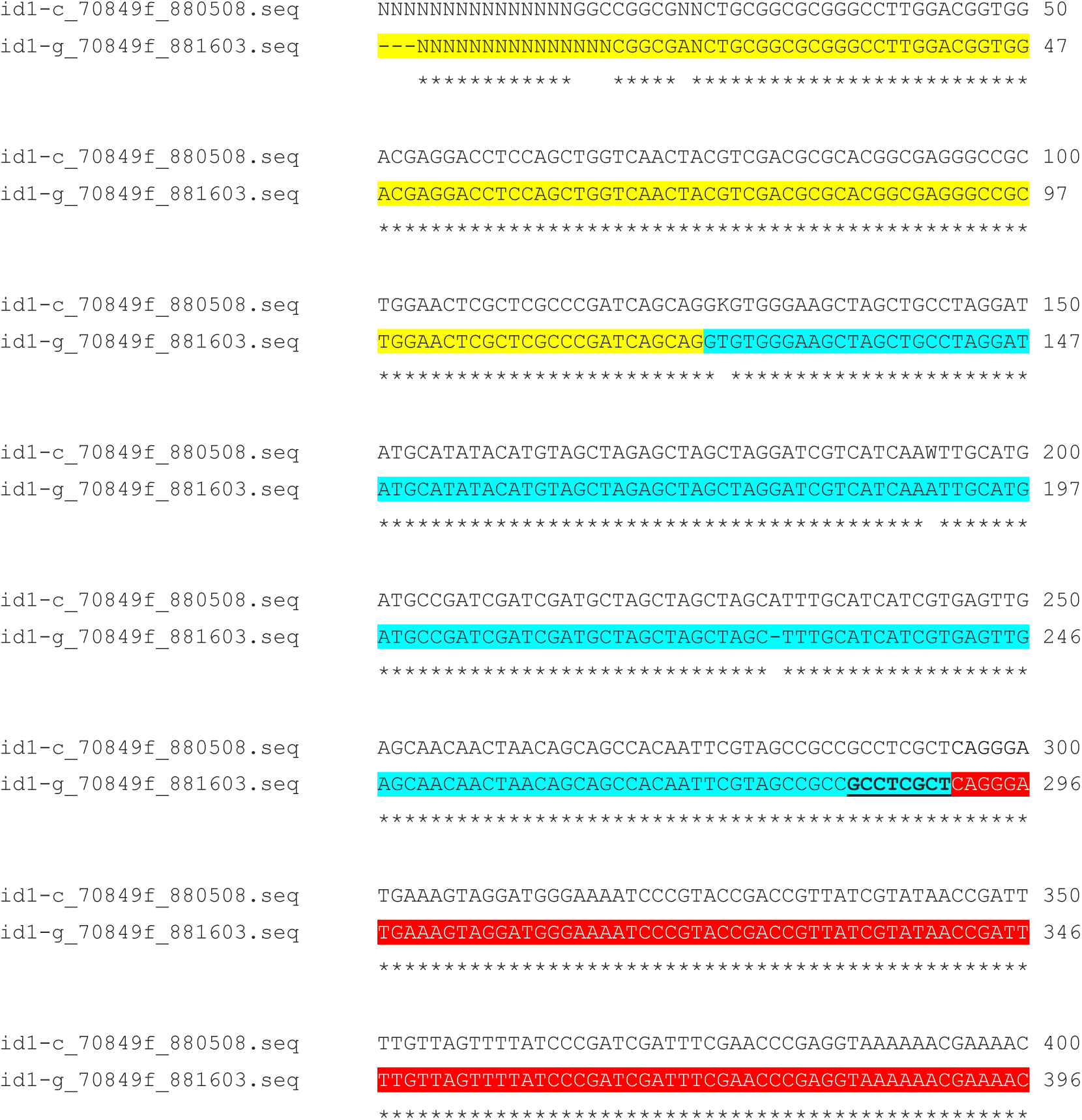

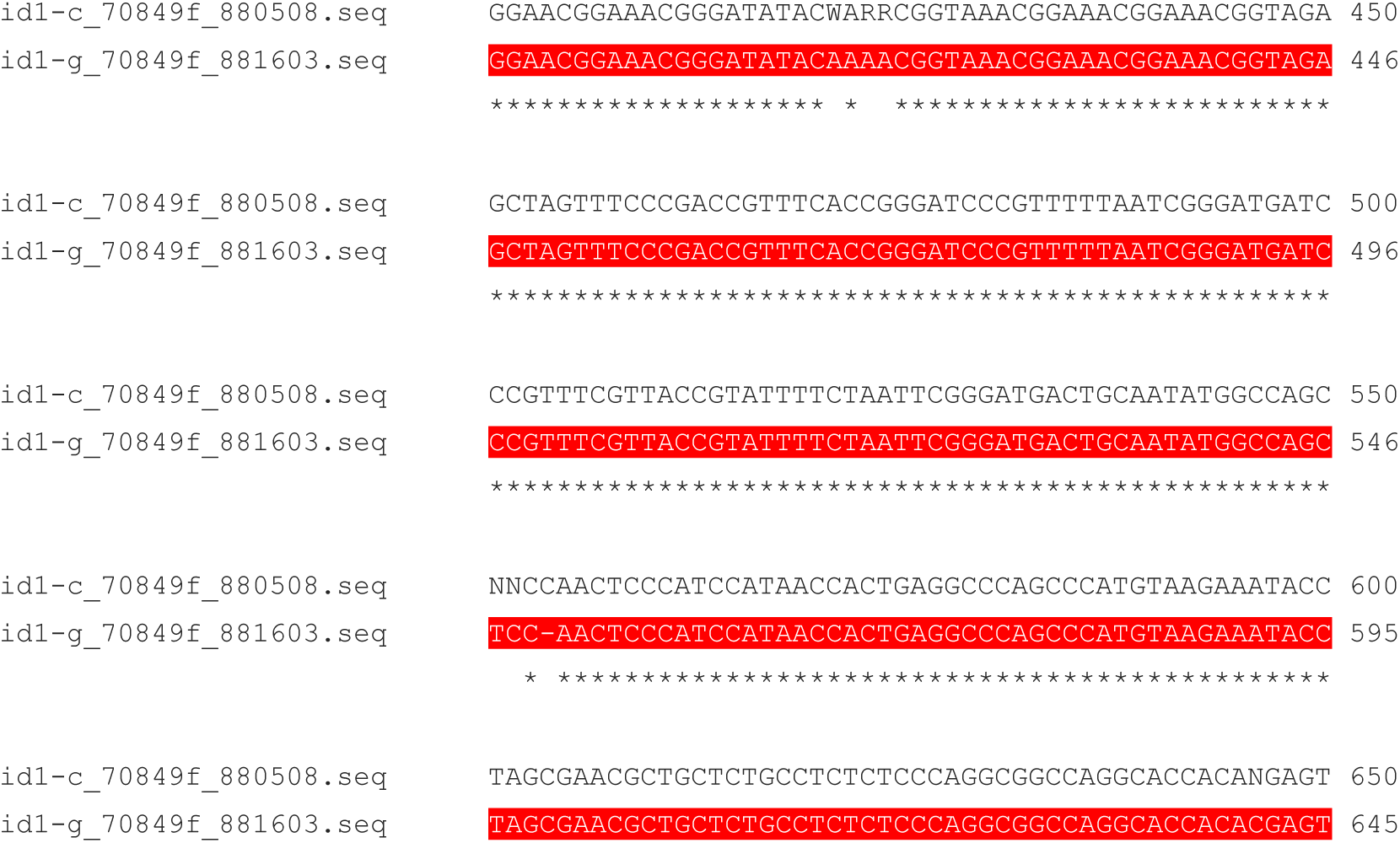

**B.** CI transcript in *p1-ww-id1* includes Intron 1 of *Ac*. Sequences were obtained from 3’ region of *p1-ww-id1* root transcript using primer Ac1568r. Upper sequence is from cDNA of chimeric transcript; lower sequence is from *Ac*.

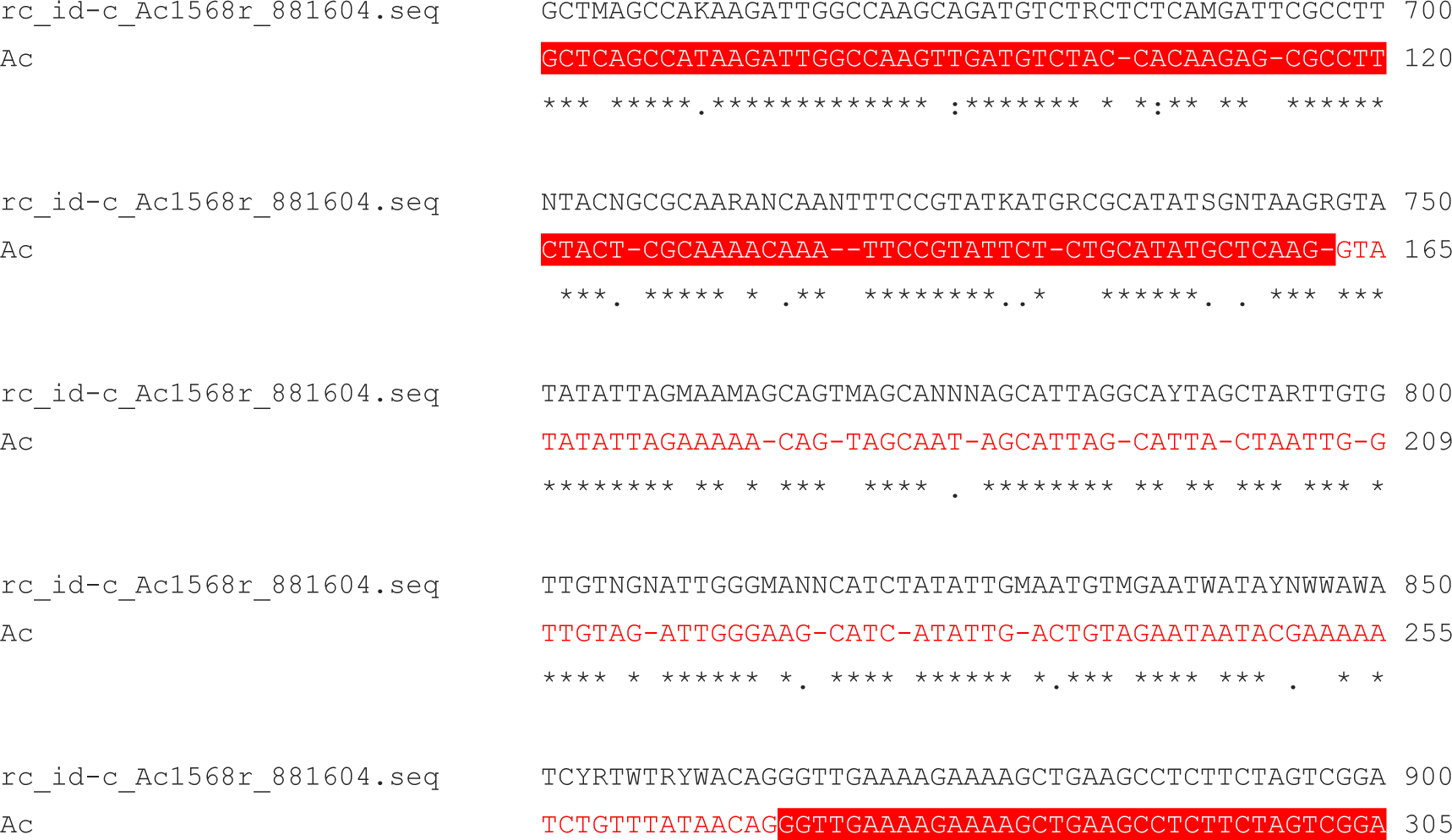

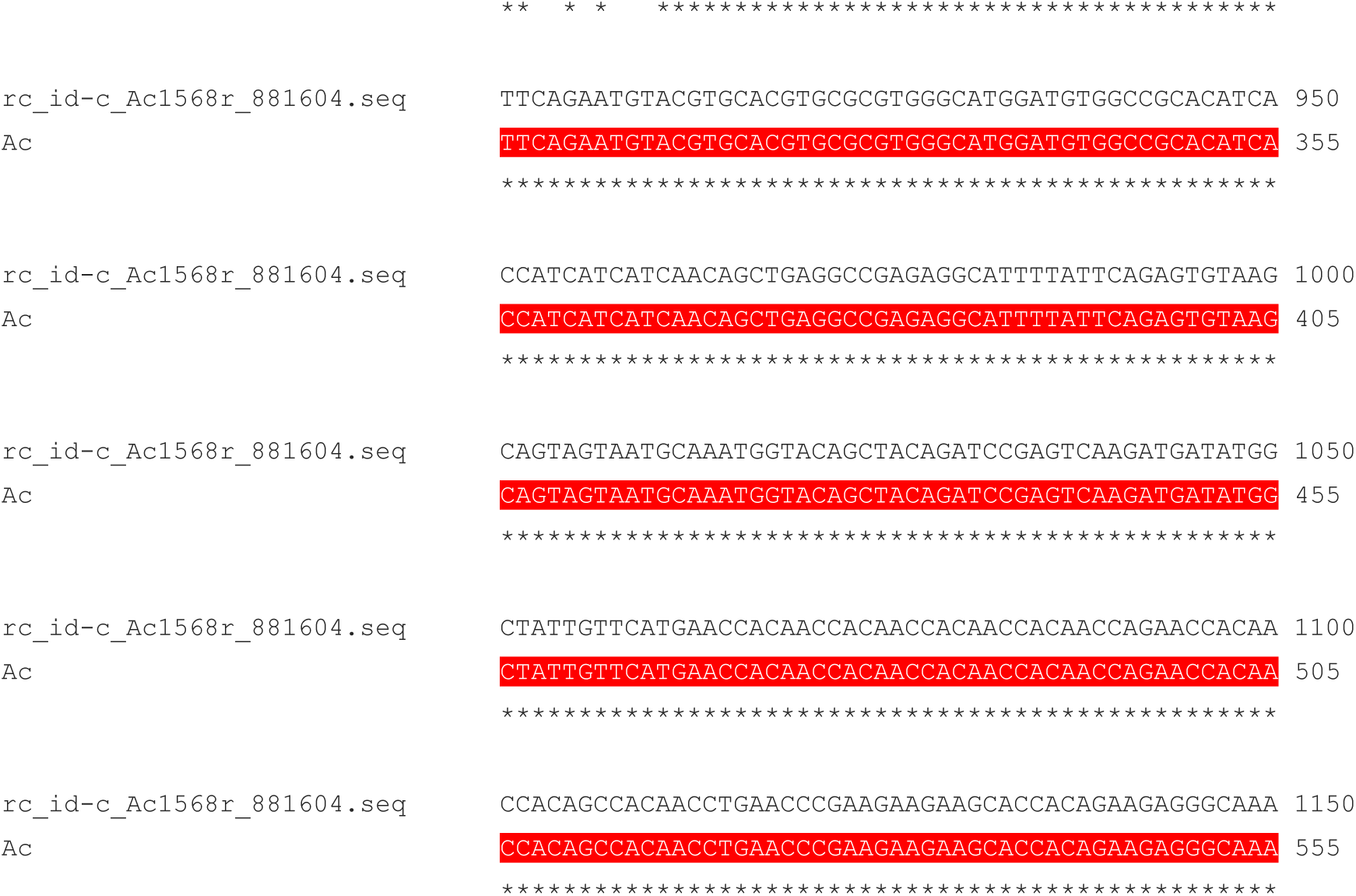

